# Functional, transcriptomic, and proteomic profiles of human primary and stem cell-derived beta cells in a state of high insulin production and increased fragility

**DOI:** 10.64898/2026.08.05.742945

**Authors:** Chieh Min Jamie Chu, Meltem E. Omur, Jasmine Maghera, Haoning Howard Cen, Alyssa Weinrauch, Sing-Young Chen, Luo Ting Helen Huang, Renata Moravcova, Jason C. Rogalski, Bhavya Sabbineni, Niki Shahraki, Samantha Mar, Cara E. Ellis, Wyeth W. Wasserman, Patrick E. MacDonald, Francis C. Lynn, James D. Johnson

**Author notes:** **Address correspondence to:** James D. Johnson, PhD., Faculty of Medicine, Department Cellular and Physiological Sciences, The University of British Columbia, Life Sciences Institute, 5358 – 2350 Health Sciences Mall, Vancouver, British Columbia, Canada V6T 1Z3, Social Media: @JimJohnsonSci.

## Abstract

Insulin production is a cardinal feature of pancreatic β cells. Studies in rodents show that β cells can switch between low and high insulin gene activity states and that elevated insulin production makes β cells more vulnerable to stresses associated with diabetes. In people, genetically elevated insulin production increases the risk of type 1 diabetes. Via effects on obesity, hyperinsulinemia contributes to the pathogenesis of type 2 diabetes. Here, we characterize β cells in low and high *INS* gene activity states sorted from primary human islets transduced with *INS*-GFP adenovirus and differentiated *INS*-EGFP knock-in embryonic stem cells (SCβ cells). We profile β cell function, protein synthesis, resilience to diabetes associated stress, single β cell transcriptomes and their co-activity networks, and purified β cell proteomes. We show that human β cells transition between distinct states. High *INS* cells have elevated maturity marker mRNAs and proteins, increased protein translation, are larger, but also more susceptible to cell death when exposed to diabetes-relevant stresses. We also catalogue thousands of differences in proteins in high *INS* stem cell-derived β cells compared directly with high *INS* primary β cells. Our study improves our understanding of the delicate balance between insulin production and β cell resilience and guides the engineering of better β cells.

**Blurb:** transcriptional, proteomic, and functional analyses of insulin gene expression states in human β cells from donor islets and stem cells

**Key findings:**

1. Functional, transcriptomic, and proteomic analyses identify similarities and differences between high and low *INS* gene activity states in primary and stem cell-derived β cells.
2. We characterize the relationship between insulin production and fragility, demonstrating that increased insulin production comes at a cost of reduced resilience to multiple stresses.
3. We report a comprehensive side-by-side proteomic analysis of purified primary and stem cell-derived β cells in the high *INS* state and identify differences in protein production and secretion machinery, providing a roadmap for making better β cells.
4. Proteomic analyses of unfolded protein response markers and cell death effectors reveal key differences in how primary and stem cell-derived β cells respond to the stress of high insulin production.

## Introduction

Pancreatic β cell dysfunction occurs early and persists throughout the course of both type 1 and type 2 diabetes. In type 1 diabetes, insulin production is paradoxically higher in those who carry risk alleles at the *INS* locus^1^ and insulin is a primary autoantigen^2^. Once disease pathogenesis starts, residual β cells must work harder. Indeed, islets from autoantibody-positive donors exhibit increased proinsulin^3,4^. Type 2 diabetes in many individuals starts with obesity^5^, which may be driven by β cell overwork and hyperinsulinemia^6,7^. Later in the pathogenesis of type 2 diabetes, β cell exhaustion and apoptosis are also associated with insulin resistance and hyperinsulinemia^8^. Rodent β cells were shown to exist in^9–12^ and transition between^13,14^ distinct low and high insulin gene activity states. Mouse cells in the elevated insulin production state were significantly more fragile when exposed to stresses associated with diabetes^13,14^. Genetically reducing insulin production in mice alleviated endoplasmic reticulum (ER) stress and allowed more β cells to enter the cell cycle^15^. These studies showed that insulin production can have direct and indirect effects on β cell fate, which may contribute to the pathogenesis of both types of diabetes. However, the functional dynamics and molecular profiles of β cell states have not been explored in human β cells, and the molecular mechanisms that govern the relationship between insulin production and β cell vulnerability are unclear.

Stem cell-derived (SC)β cell transplantation holds promise for replacing the ∼80-90% of insulin-producing cells destroyed by autoimmunity in type 1 diabetes^16^. Several hurdles must be cleared before this can become a common clinical reality^17^. First, SCβ cells will have to be protected from alloimmunity and autoimmunity via encapsulation, and/or genetic engineering with immune-evasive proteins, and/or improvement of iPSC-based protocols to make cells more robust. Second, more stress-resistant SCβ cells are required to better survive the initial post-transplant period and last longer than grafts are currently predicted. Third, SCβ cells still require better regulated insulin production and secretion at baseline and in response to nutrient stimulation. While protocols for producing SCβ cells have improved over the past years and it is now possible to consistently make glucose-responsive cells, there is not yet reproducible evidence that they have similar insulin-release dynamics when compared to high-quality primary human islets^17–19^. Clearing these hurdles will require detailed understanding of the mechanisms controlling and linking insulin production states and cell survival in SCβ cells. Side-by-side, deep profiling of primary and stem cell-derived β cells provides a unique opportunity to map the remaining deficiencies in *in vitro* differentiation protocols.

In the present study, we investigated insulin production states in primary human β cells and differentiated SCβ cells with function and survival assays, scRNAseq, and mass-spec proteomics on sorted cells. This comprehensive comparison between low and high *INS* production states within the same cell cultures define key characteristics of β cells in their high working state that may shape their susceptibility to stresses associated with diabetes progression and following cell replacement therapy.

## Methods

### Culture and INS-GFP adenoviral transduction of human primary β cells

Islets were isolated from the pancreata of cadaveric human donors (Table S1) at the Alberta Diabetes Institute IsletCore (www.isletcore.ca). All families of organ donors provided written informed consent for use in research. T2D diagnosis was family-declared at the time of organ donation. Islet isolation was performed as described^20^. Islets were used with approval of the Human Research Ethics Board at the University of Alberta (Pro00013094; Pro00001754) and the University of British Columbia Clinical Research Ethics Board (H13-01865). Islets were shipped overnight in CMRL media (Thermo Fisher Scientific, Waltham, USA) and further purified after arrival by handpicking under a stereomicroscope. Islets were cultured in RPMI 1640 medium (Thermo Fisher Scientific, Cat#, 11879-020) supplemented with 5.5 mmol glucose, 10% fetal bovine serum (FBS), and 100 units/mL penicillin/streptomycin in 10 cm non-adhesive petri dishes (Thermo Fisher Scientific, Cat # FB0875713) for 24-72 hours prior to experiments to allow for recovery from shipment. To label insulin-producing cells with GFP we used an adenovirus with GFP driven by a rat *Ins2* promoter^9^ (gift from T. Kieffer). Islets in 15 mL tubes were treated with Accutase (STEMCELL Technologies, Canada) for 8 minutes, with gentle pipetting to disperse the cells. Cells were incubated with 100 MOI adenovirus for 1 hour with gentle shaking. Cells were then seeded onto 12-well plates and allowed to recover for 48 hours before experiments.

### Differentiation and dispersion of stem cell derived spheroids

Human embryonic stem cells (WiCell WA01 line) that contain eGFP knocked-in and added-on downstream of the insulin coding sequence were differentiated to stem cell-derived β-like (SCβ) cells using a 6-stage protocol as described^21^. After the initial 21 days of differentiation, SCβ cells were cultured in CMRL with 5.6 mM glucose (Thermo Fisher Scientific) containing 1% fatty acid-free BSA (Proliant), 1:100 Glutamax, 1:100 NEAA, 1 mM pyruvate, 10 mM HEPES, 1:100 ITS (Thermo Fisher Scientific), 10 μg/ml of heparin sulfate, 1 mM N-acetyl cysteine, 10 μM zinc sulfate, 1.75 μL 2-mercaptoethanol, 2 μM T3 and 0.155 mM ascorbic acid (Sigma Aldrich) until Day 27. Between Day 27-35, aggregates were grown in CMRL with 5.6 mM glucose containing 1% fatty acid-free BSA, 1:100 Glutamax, 1:100 NEAA, 1 mM Pyruvate, 10 mM HEPES, 1:100 ITS, 10 μg/ml heparin sulfate, 1 mM N-acetyl cysteine, 10 μM zinc sulfate, 1.75 μL 2-mercaptoethanol, 10 nM T3, 1:2000 Trace elements A (Cellgro), 1:2000 trace elements B (Cellgro), 1:2000 Lipid Concentrate (Thermo Fisher Scientific) and 0.5 µM ZM447439 (Selleckchem)^18^. In some studies, we used a variant of this protocol wherein WIKI4 is added at stage 4, described elsewhere^22^. We define the age of the SCβ cells as the total number of days in culture through all stages. For dispersion into individual SC β cells, spheroids were hand-picked and washed 2 times with PBS. Spheroids were then immersed for 8 minutes in Accutase (STEMCELL technologies, Vancouver, Canada), with gentle flicking every 2 minutes, 4x dilution with PBS to halt Accutase activity, then centrifugation for 5 min at 200 x g. Cells were re-suspended in media with 10 uM ROCK inhibitor Y-27632 (STEMCELL Technologies, Vancouver, Canada).

### Fluorescence-activated cell sorting

Dispersed cells were filtered into 5 ml polypropylene tubes and sorted at the Life Sciences Institute core facility using a Cytopeia Influx (Becton Dickinson, Franklin Lakes, NJ, USA) with excitation at 488 nm (530/40 emission) and 561 nm (610/20 emission). In cases where we performed a reaggregation step, SCβ spheroids were dispersed as above and purified using fluorescence-activated cell sorting (FACS) into Aggrewell 800 plates (STEMCELL Technologies, Vancouver, Canada) to generate clusters containing 1,000 cells/cluster. After 60 hours, the reaggregated clusters were transferred into 6-well plates containing Stage 7 media.

### Live cell imaging

For 3D live imaging, intact day 35 stem cell-derived spheroids were incubated overnight in CMRL media (5.6 mM glucose, 1% fatty acid-free BSA, 1:100 Glutamax, 1:100 NEAA, 1 mM pyruvate, 10 mM HEPES, 1:100 ITS, 10 μg/ml heparin sulfate, 1 mM N-acetyl cysteine, 10 μM zinc sulfate, 1.75 μL 2-mercaptoethanol, 10 nM T3, 1:2000 trace elements A, 1:2000 trace elements B, 1:2000 lipid concentrate and 0.5 µM ZM447439) with 0.05 μg/mL Hoechst 33342 and 0.5 μg/mL propidium iodide. Cells were imaged in 8 well µ-Slides (Ibidi) in fresh culture media using a LEICA THUNDER Live Cell & 3D microscope at 25x water objective (numerical aperture 0.95). Image processing, including smoothing and background subtraction, was performed using Imaris analysis software. The Labkit FIJI module facilitated image labeling and pixel classification for identification of GFP fluorescence in individual β cells. Nearest neighbor analysis was done using Imaris Python XTensions. Distance-to-core analysis was conducted on Imaris, using a reference frame positioned at the islet’s center to measure distance of each cell from the origin reference frame.

To image cell death^23^, cells were seeded (8000 cells/well for SCβ; 4000 cells/well for primary) onto 384-well imaging plates (PerkinElmer, Waltham, United States) coated with Cultrex (R&D systems, Minneapolis, United States). The next day, cells were washed with PBS then cultured in media with 0.05 μg/mL Hoechst 33342 and 0.5 μg/mL propidium iodide starting 2 h before imaging using an ImageXpress Micro Confocal environmentally-controlled, robotic imaging system (Molecular Devices, San Jose, United States). Images were acquired with 10x air objective (0.3 numerical aperture) at 2 h intervals up to 96 h with the following exposures - 359 nm for 110 ms; 491 nm for 15 ms; 561 nm for 75 ms. GFP tracking in individual cells was performed using custom MetaXpress scripts and R; high and low *INS* activity states were identified using model based clustering^24^.

### Static glucose stimulated insulin secretion on sorted cells

Glucose-stimulated insulin and glucagon secretion were assessed by seeding FACS purified cells (10,000 cells per well) into a 384 well imaging plate (PerkinElmer, Waltham, United States) that was pre-coated with Cultrex (R&D systems, Minneapolis, United States). Cells were allowed to adhere for 24 hours in culture media (CMRL with 5.6 mM glucose containing 1% fatty acid-free BSA, 1:100 Glutamax, 1:100 NEAA, 1 mM Pyruvate, 10 mM HEPES, 1:100 ITS, 10 μg/ml heparin sulfate, 1 mM N-acetyl cysteine, 10 μM zinc sulfate, 1.75 μL 2-mercaptoethanol, 10 nM T3, 1:2000 Trace elements A, 1:2000 trace elements B, 1:2000 Lipid Concentrate and 0.5 µM ZM447439). Cells were washed Krebs–Ringer Buffer (KRB; 129 mM NaCl, 4.8 mM KCl, 1.2 mM MgSO4, 1.2 mM KH2PO4, 2.5 mM CaCl2, 5 mM NaHCO3, 10 mM HEPES, 0.5% bovine serum albumin) containing 3 mM glucose then pre-incubated for 4 h in 3 mM glucose KRB. Cells were incubated in KRB with 3 mM glucose then 20 mM glucose for 45 min each with addition of leucine (5 mM) and palmitate/oleate (1.5 mM). Supernatant was collected after each stimulation, and cell loss after treatments and media collection was assessed using a robotic imaging system (Molecular Devices, San Jose, United States). Insulin was assessed by Human Insulin Chemiluminescent ELISA (ALPCO: 80-INSMR) and measured on a Spark plate reader (TECAN). Glucagon was assessed using MSD assays (Mesoscale Discovery, Maryland, United States).

### Protein synthesis assay

One day after dispersion, FACS purified primary human β cells and SCβ cells were seeded into an optical 384-well plate (Perkin Elmer, Waltham, United States) at a density of 8,000 cells per well in culture media (CMRL with 5.6 mM glucose containing 1% fatty acid-free BSA, 1:100 Glutamax, 1:100 NEAA, 1 mM Pyruvate, 10 mM HEPES, 1:100 ITS, 10 μg/ml heparin sulfate, 1 mM N-acetyl cysteine, 10 μM zinc sulfate, 1.75 μL 2-mercaptoethanol, 10 nM T3, 1:2000 Trace elements A, 1:2000 trace elements B, 1:2000 Lipid Concentrate and 0.5 µM ZM447439). Treatments were applied 3 hours after seeding. After 24 hours of incubation, fresh culture media was applied, then supplemented with 20 μM O-propargyl-puromycin (Invitrogen, Waltham, United States). The assay was performed according to instructions provided by the manufacturer. Cells were imaged at 10x, numerical aperture 0.3, with an ImageXpress^Micro^ high-content imager and analyzed with MetaXpress to quantify the integrated staining intensity of OPP-Alexa Fluor 594 in cells identified by NuclearMask Blue Stain. All data analyzed in these assays were normalized to cell size.

### Single cell RNAseq

To define the relationship between single cell *INS* mRNA levels and single-cell function, we reanalyzed a previously published patchSeq data were obtained from the NCBI Gene Expression Omnibus (GEO) and Sequence Read Archive (SRA) under accession numbers GSE124742, GSE164875 and GSE270484; at PancDB (https://hpap.pmacs.upenn.edu/); and at Human Cell Atlas Data Explorer under project label CryoPancreaticIsletCellPatchSeq. Electrophysiology correlations used the same within-study meta framework described above, pooled across 50 imputations to handle missing electrophysiology data on the z-scale, with 95% CIs back-transformed to ρ.

To compare the transcriptomic profiles of *INS* mRNA low and high β cells, we reanalyzed a large compilation of scRNAseq from 60,663 primary human β cells from 69 non-diabetic donors is detailed elsewhere^25^. Here, cells were reclassified into None, Low, and High *INS* expression groups using Gaussian mixture modeling (previously published data available from the NCBI Gene Expression Omnibus (GEO) and Sequence Read Archive (SRA) under accession numbers GSE81076, GSE81547, GSE81608, GSE83139, GSE84133, GSE85241, GSE86469, GSE154126, GSE269204; from the European Molecular Biology Laboratory European Biology Institute (EMBL-EBI) under accession number E-MTAB-5061; from the Human Pancreas Analysis Program at PancDB up to donor HPAP-122). Differential expression and downstream GSEA comparing the low and high *INS* expression groups was performed using the glmGamPoi pseudobulk workflow, with *INS* expression came from the RNA/counts layer (raw counts). For each gene, we also computed Spearman correlations within each donor/study with *INS* expression (from the RNA/data layer, log-normalized counts), then combined donors via a random-effects meta-analysis on the Fisher-z scale with Hartung–Knapp–Sidik–Jonkman confidence intervals (RE+HKSJ). To calibrate significance while respecting donor structure, we additionally ran donor-blocked permutations: within each donor, cells were binned into Q=10 equal-count *INS* quantiles; we correlated bin means vs 1…Q and pooled donors (fixed-effects on z). We did B=1000 within-donor permutations of bin order to obtain empirical two-sided p-values and BH FDR (FDR_perm). Figures show RE+HKSJ ρ and CI; point sizes reflect FDR_perm.

For SCβ cell patchSeq analysis, multi-omics sequenced files were processed and analyzed using Cell Ranger version 8. Genes were mapped and referenced using the human reference genome GRCh38. RNA modality from integrated dataset is used for downstream analysis. Low-quality cells were removed by filtering out cells with low RNA counts (nCount_RNA <300). Cell type annotation was performed manually by the expression of specific markers. SCβ cells were subsetted and categorized based on mean expression levels of insulin and EGFP. Of the SCβ 344 cells exhibiting higher expression levels of both *INS* and *egfp* than the mean values were annotated as *INS*(EGFP)^HIGH^, while 482 cells with expression levels lower than the mean values for both *INS* and *egfp* were classified as *INS*(EGFP)^LOW^. ‘CorrelatePairs’ function in R is used to calculate gene pairs correlation. Pheatmap package and ‘DoHeatmap’ from Seurat package were used for plotting heatmaps. ‘DotPlot’ and ‘VlnPlt’ functions from the Seurat package are used to plot expression of genes. Pathway analysis was performed with ‘EnrichGo’ function with ‘org.Hs.eg.db’ as an organism database from ClusterProfiler 4.2.2 and org.Hs.eg.db 3.14.0 packages. ‘barplot’ function in R is used to visualize pathway results.

A larger SCβ single cell dataset was generated using 10X Chromium Next GEM Single Cell Multiome ATAC + Gene Expression and Chromium Nuclei Isolation (CG000505 Rev A) kits according to the manufacturer’s protocol. Multi-omics sequenced files were processed and analyzed using Cell Ranger version 6.1.1. Genes were mapped and referenced using the human reference genome GRCh38. Datasets were analyzed using Seurat 4.3.0 in R 4.1.1 within Jupyter Notebook 7.0.2. RNA modality from integrated dataset was used for downstream analysis. Low-quality cells were removed by filtering out cells with low RNA counts (nCount_RNA <300). Cell type annotation was performed manually by the expression of specific markers. SCβ cells were subsetted and categorized based on mean expression levels of insulin and EGFP. 344 cells exhibiting higher expression levels of both *INS* and *egfp* than the mean values were annotated as *INS*(EGFP)^HIGH^, while 482 cells with expression levels lower than the mean values for both *INS* and *egfp* were classified as *INS*(EGFP)^LOW^. Cells that do not fall into either of these categories are excluded from analysis. ‘CorrelatePairs’ function in R 4.1.1 is used to calculate gene pairs correlation. Pheatmap package and ‘DoHeatmap’ from Seurat 4.3.0 package were used for plotting heatmaps. ‘DotPlot’ and ‘VlnPlt’ functions from the Seurat 4.3.0 package were used to plot expression of genes. Pathway analysis was performed with ‘EnrichGo’ function with ‘org.Hs.eg.db’ as an organism database from ClusterProfiler 4.2.2 and org.Hs.eg.db 3.14.0 packages. ‘barplot’ function in R 4.1.1 is used to visualize pathway results.

### SCENIC analysis

Gene regulatory network (GRN) inference was performed using pySCENIC^26^ separately on the quality-controlled primary human β cell and SCβ scRNA-seq datasets. For primary β cells, the GRN inference step (pyscenic grn) was run 40–100 times; transcription factor-target edges present in ≥80% of runs were retained and edge importance scores were averaged to yield a consensus GRN, which was passed to the cisTarget (ctx) and AUCell steps to generate regulon definitions and per-cell regulon activity scores (AUC values). For SCβ cells (all cell populations), the GRN inference and cisTarget steps were run 20 times; interaction pairs present in ≥80% of runs were retained and used as input for AUCell calculation.

For co-activity network analyses, we constructed regulon co-activity networks separately for beta cell and SCβ subsets with high vs. low *INS* expression using SCENIC AUC scores. For each group (*INS*-high and *INS*-low), we performed 100 bootstrap iterations, sampling cells with replacement to capture variability within each group and subsampling from the *INS*-high group for a balanced design. In each iteration, we computed pairwise rank-based correlations (Spearman’s ρ) between regulon AUC scores across the sampled cells. The resulting correlation matrices were thresholded (|ρ| > 0.6 for primary β cells, |ρ| > 0.45 for SCβs) to retain only strong co-activity signals, and converted into undirected, weighted igraph objects: nodes represent transcription factors (regulons), while edges represent strong positive or negative co-activity based on ranked expression similarity. Edge weights were set to the absolute correlation coefficient. For each bootstrap network, we calculated the following graph topology metrics: 1) Number of nodes and edges 2) Network density and mean degree 3) Global clustering coefficient 4) Degree skewness 5) Louvain modularity 6) Diameter and mean path length of the largest connected component. Metrics were summarized and compared across groups using faceted boxplots and summary statistics.

### Mass spec proteomics

Cells (89,000-500,000) were diluted with cold ultrapure water and lysed with 50% trifluoroethanol (TFE), chilled for 10 min, vortexed for 1 min, sonicated on ice for 10 min, then boiled at 95°C for 10 min; this cycle was repeated once. pH was adjusted to 8-8.5 using 1M Tris-HCl (final concentration 100 mM). Total protein was estimated at 0.25 ng/cell. Reduction and alkylation used tris(2-carboxyethyl)phosphine hydrochloride/chloroacetamide (TCEP/CAA) was added to a final concentration of 10 mM/40 mM and 5 min at 70°C. Samples were diluted with 50 mM ammonium bicarbonate. Proteins were digested with trypsin overnight at 37°C (1:100 enzyme:protein, w/w), followed by a second digestion for 2-3 hours at 37°C (1:125, w/w) before quenching with 10% formic acid. Peptides were desalted using C18 STAGE-tips^27^ and reconstituted in 0.5% acetonitrile, 0.1% formic acid. Peptide concentration was measured by absorbance at 205 nm (baseline correction at 340 nm) using NanoDrop (ThermoFisher, Waltham, USA). Up to 200 ng peptides were injected and separated on NanoElute UHPLC system (Bruker Daltonics, Billerica, United States) with Aurora Series Gen2 (CSI) analytical column (Ion Opticks, Parkville, Australia), heated to 50°C, coupled to timsTOF Ultra 2 (Bruker Daltonics, Billerica, USA) operated at 0.3 μL/min and 7°C. Buffer A consisted of 0.1% formic acid and 0.5 % acetonitrile in water, and buffer B consisted of 0.1% formic acid in 99.5% acetonitrile. The analytical column was conditioned with 4 column volumes of buffer A prior to each run. Peptides were eluted using 60-min gradient from 2% B to 12% B over 30 min, then to 33% B by 60 min, ramped to 95% B over 0.5 min, and held at 95% B for 7.72 min. Capillary voltage was set to 1600V, drying gas flow to 3 L/min, and drying temperature to 200°C. Each MS1 scan was followed by 20 variable PASEF ramps, containing 60 MS/MS isolation windows covering m/z 350.7– 1250.6. Ion mobility range (1/k0) was set to 0.66 – 1.42 V·s/cm2, 50 ms ramp and accumulation time (100% duty cycle) at a ramp rate of 17.80 Hz; resulting in a total cycle time of 1.18s. Collision energy was ramped linearly as a function of mobility from 20 eV at 1/k0 = 0.6 V·s/cm2 to 59 eV at 1/k0 = 1.60 V·s/cm2. Mass-to-charge and ion mobility were calibrated using three Agilent ESI-Low Tuning Mix ions (m/z [Th], 1/k0 [Th]: 622.0290, 0.9915; 922.0098, 1.1986; 1221.9906, 1.3934), with mass accuracy typically within 4 ppm and not exceeding 7 ppm (timsControl v. 6.0.6 and HyStar 6.3). Two spectral libraries were generated: one for sorted stem cells and one for sorted human islets. For each, 12 or 20 samples respectively were randomly selected, and equal amounts were pooled to create a representative sample. The pooled sample was injected four times and separated using narrow gas-phase fractionation windows, with isolation windows reduced to one-quarter of those used in sample acquisition.

DIA data were searched library-free using DIA-NN (v2.1.0 Academia^28^) against a *Homo sapiens* database (reviewed sequences only; 20390 entries from Uniprot Nov 03, 2025) supplemented with insulin chain A and B, proinsulin, proglucagon, IAPP, green fluorescent protein, and red fluorescent protein drFP583, and common contaminants (211 entries). Searching used trypsin/P specificity with 1 missed cleavage, N-terminal M excision and cysteine carbamidomethylation as fixed modification. Peptide length restricted to 7-30 residues, precursor charge to 2-4, precursor m/z to 300-1200, and fragment ion m/z to 200-1800. Precursor FDR was 1%, with MS1 accuracy and MS2 accuracy set to 15, and calibration accuracy and scan window set to 0. Heuristic protein inference, isotopologues usage, match between run (MBR), and no shared spectra were all enabled. Gene-level protein interference was chosen with double-pass neural network classification. Quantification was performed using Robust LC (high precision) mode, with cross-run normalization disabled. Experimental samples were searched using DIA-NN with the same parameters, with the addition of a previously generated spectral library. Non-normalized data was then processed with PRONE in R using the Normics median normalization method^29^. Proteins which were in less than 80% of at least one cell population per comparison were removed. Paired t-tests in Perseus were adjusted for multiple comparisons.

### Statistics and data visualization

Statistics and data representation for nearest neighbor analyses, distance to core analyses, and translation assays, were conducted using GraphPad Prism 10 (Graphpad Software, San Diego, USA). Statistics and data representation for live cell imaging experiments utilized custom R scripts. Student’s t-test, one-way ANOVAs, and two-way ANOVAs were used for parametric data analysis as indicated in figure legends. For analyses of mass-spec proteomics, we utilized perseus software and performed student’s t-test adjusted for multiple comparisons. For all statistical analyses, differences were considered significant if the *p* value or adjusted *p* value was less than 0.05. Error bars represent ± standard error of the mean (SEM).

## Results

### INS production states in human β cells

To image insulin production state dynamics in living human β cells, we infected dispersed islet cells with adenovirus carrying a GFP reporter under control of an insulin promoter (Fig. 1A, Supplemental Fig. 1A). We previously validated that similar promoter reporters faithfully report insulin production in mouse and human β cells^11,12^. Here, we used FACS to categorize GFP+ cells into *INS*(GFP)^LOW^ and *INS*(GFP)^HIGH^ gene activity (Fig. 1B), consistent with the low and high insulin production states we have previously reported in *Ins2*^GFP^ knock-in mouse β cells^13,14^. Long-term imaging showed that GFP fluorescence, reflecting *INS* gene activity, was glucose-responsive with *INS*(GFP)^HIGH^ cells reacting faster and having a greater increase in GFP compared to *INS*(GFP)^LOW^ cells (Fig. 1C,D; Supplemental Fig. 1B). There was a tendency for increased *INS*(GFP)^LOW^ to *INS*(GFP)^HIGH^ transitions and reduced *INS*(GFP)^HIGH^ to *INS*(GFP)^LOW^ transitions with higher concentrations of glucose (Fig. 1C,D). Most cell state transitions were *INS*(GFP)^HIGH^ to *INS*(GFP)^LOW^ transitions (Supplemental Fig. 1C), similar to primary *Ins2*^GFP^ mouse β cells^13,24^. Inducing ER stress with thapsigargin (Tg) drove *INS*(GFP)^HIGH^ to *INS*(GFP)^LOW^ transitions and decreased in *INS*(GFP)^LOW^ to *INS*(GFP)^HIGH^ transitions (Supplemental Fig. 1D). These experiments illustrate the potential insulin production states to be dynamic and responsive to nutrients and stress in human β cells.

**Figure 1.**
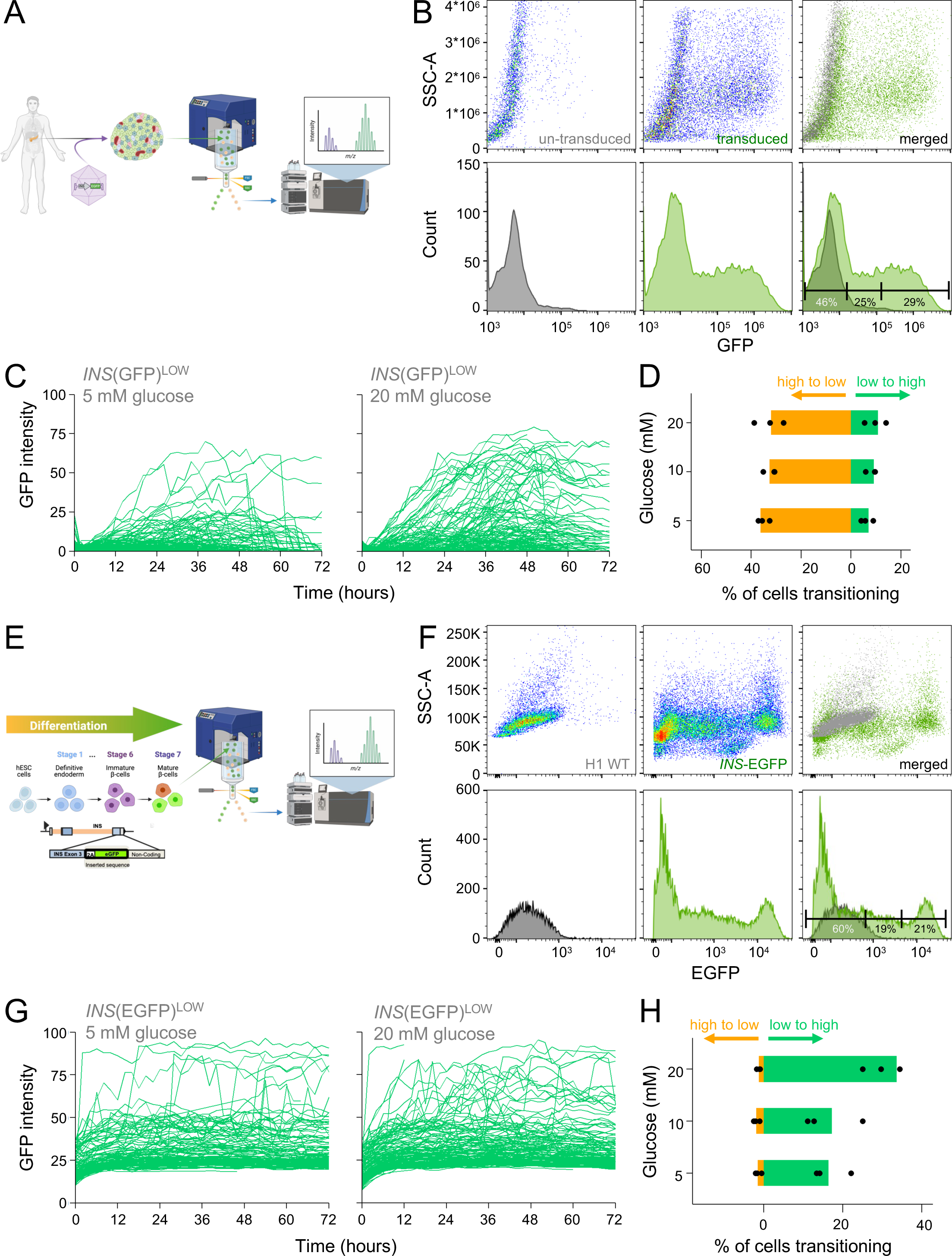
*INS* production states in human insulin producing cells. **(A)** Schematic and experiment layout of human primary β cells transduced with an *INS*-GFP adenovirus. **(B)** Identification of a bimodal distribution in *INS* production by FACS in primary β cells. **(C)** Example of tracking the GFP of individual primary β cells over time. Each line represents a cell. **(D)** Cell state transitions in primary β cells between *INS*(GFP)^HIGH^ and *INS*(GFP)^LOW^ states in various culture conditions (n=3 human islet donors) **(E)** Schematic and experiment layout of the *INS*-EGFP SCβ cells. **(F)** Identification of a bimodal distribution in *INS* production by FACS in SCβ cells. **(G)** Example of tracking the GFP of individual SCβ cells over time. Each line represents a cell. **(H)** Cell state transitions over 48 hours in SCβ cells between *INS*(EGFP)^HIGH^ and *INS*(EGFP)^LOW^ states in media containing ITS but no thapsigargin conditions (n=3 wells).

As a complementary human β cell model, we studied SCβ cells with *INS*-EGFP knock-in at Day 35 and Day 50 of a 7-stage protocol^19^ (Fig. 1E). After controlling for autofluorescence using H1 cells, FACS showed that EGFP+ cells can be clearly categorized into *INS*(EGFP)^LOW^ and *INS*(EGFP)^HIGH^ states (Fig. 1F). Similar to the primary β cells, *INS*(EGFP)^HIGH^ cells had increased granularity compared with *INS*(EGFP)^LOW^ cells (Fig. 1F). The two populations were also found in Day 50 SCβ cells, as well as in cells with WIKI4 added at stage 4^22^ (Supplemental Fig. 2A,B). Addition of WIKI4 at stage 4 (protocol B)^22^ increased the percentage of cells in the *INS*(GFP)^LOW^ state, while not affecting the proportion of cells in the *INS*(GFP)^HIGH^ state (Supplemental Fig. 2C). *INS* gene activity in intact stem cell-derived spheroids also showed a bimodal distribution (Supplemental Fig. 3A,B). Nearest neighbor and distance to core analyses found that cells in the same *INS* activity state tended to physically cluster closer to one another, and that cells in the *INS*(EGFP)^HIGH^ state tended to be closer to the core of the spheroid (Supplemental Fig. 3C, D). Together, these data show that SCβ cells can exist in low and high INS gene activity states, like primary mouse and human β cells. Long term live-cell imaging revealed that SCβ cells could transition between the *INS*(EGFP)^HIGH^ and *INS*(EGFP)^LOW^ expression states; interestingly, very few cells transitioned from the *INS*(EGFP)^HIGH^ to the *INS*(EGFP)^LOW^ state (Fig. 1G, H). The majority of cells showed an initial rise in the first 12 hours of imaging conditions (Fig. 1G, Supplemental Fig. 3E). The number of cells transitioning from *INS*(EGFP)^HIGH^ to the *INS*(EGFP)^LOW^ only started to increase with longer culture conditions (Supplemental Fig. 3F). Similar to primary β cells, higher glucose concentrations increased the number of cells transitioning from *INS*(EGFP)^LOW^ to *INS*(EGFP)^HIGH^ (Fig. 1H). Contrary to the primary β cells, Tg only reduced the number of cells transitioning from *INS*(EGFP)^LOW^ to the *INS*(EGFP)^HIGH^, while not effecting the number of cells transitioning from *INS*(EGFP)^HIGH^ to the *INS*(EGFP)^LOW^(Supplemental Fig. 3G). We also investigated the stability of the two cell states by FACS purifying and reaggregating cells from the two cell states into spheroids and analyzing their compositions (Supplemental Fig. 4A, B). Interestingly, *INS*(EGFP)^HIGH^ cells formed larger reaggregates than *INS*(EGFP)^LOW^ cells (Supplemental Fig. 4A). After four days of reaggregation, plus an additional 2 days of rest, many cells in the *INS*(EGFP)^HIGH^ reaggregates transitioned to *INS*(EGFP)^LOW^ state (Supplemental Fig. 4B). In contrast, few cells from the *INS*(EGFP)^LOW^ state transitioned to *INS*(EGFP)^HIGH^ (Supplemental Fig. 4B). These results show that cell state transitions can also occur in SCβ cells, but in a manner that is distinct from primary mouse and human β cells.

### Functional characterization of human primary β cells in low and high INS states

We next characterized how *INS* activity states relate to the physiological functions of primary human β cells. FACS forward scatter indicated that primary β cells in the *INS*(GFP)^HIGH^ state were larger than cells in the *INS*(GFP)^LOW^ state (Fig. 2A). FACS side scatter (often used as an indication of granularity) was also found to be increased in the *INS*(GFP)^HIGH^ state (Fig. 2B). Adherent *INS*(GFP)^HIGH^ cells had significantly greater average area than *INS*(GFP)^LOW^ cells in live imaging experiments (Fig. 2C). Increased cell size is often a result of elevated protein synthesis. It is known that β cells rapidly expand when translating *INS* mRNA in response to metabolic demand^30^. Using O-propargyl-puromycin (OPP) to assess translational capacity and normalizing to cell size, we found cells in the *INS*(GFP)^HIGH^ state had significantly more protein synthesis compared to cells in the *INS*(GFP)^LOW^ state across all conditions tested (Fig. 2D). Similar to our live cell imaging analyses, addition of Tg was found to increase the number of cells in the *INS*(GFP)^LOW^ state (Fig. 2E). Increased protein translation, especially of insulin, may lead to ER stress^15^ and programmed cell death. Image-based survival assays^23^ with various concentrations of glucose (5, 10, 20 mM) and thapsigargin (1, 10 μM) revealed that β cells in the *INS*(GFP)^HIGH^ state were more fragile than those in the *INS*(GFP)^LOW^ state (Fig. 2F, G). These results demonstrate that primary human β cells with high insulin production are larger with higher granularity, possess higher translational capacity, but that this comes at the cost of increased fragility in all conditions tested.

**Figure 2.**
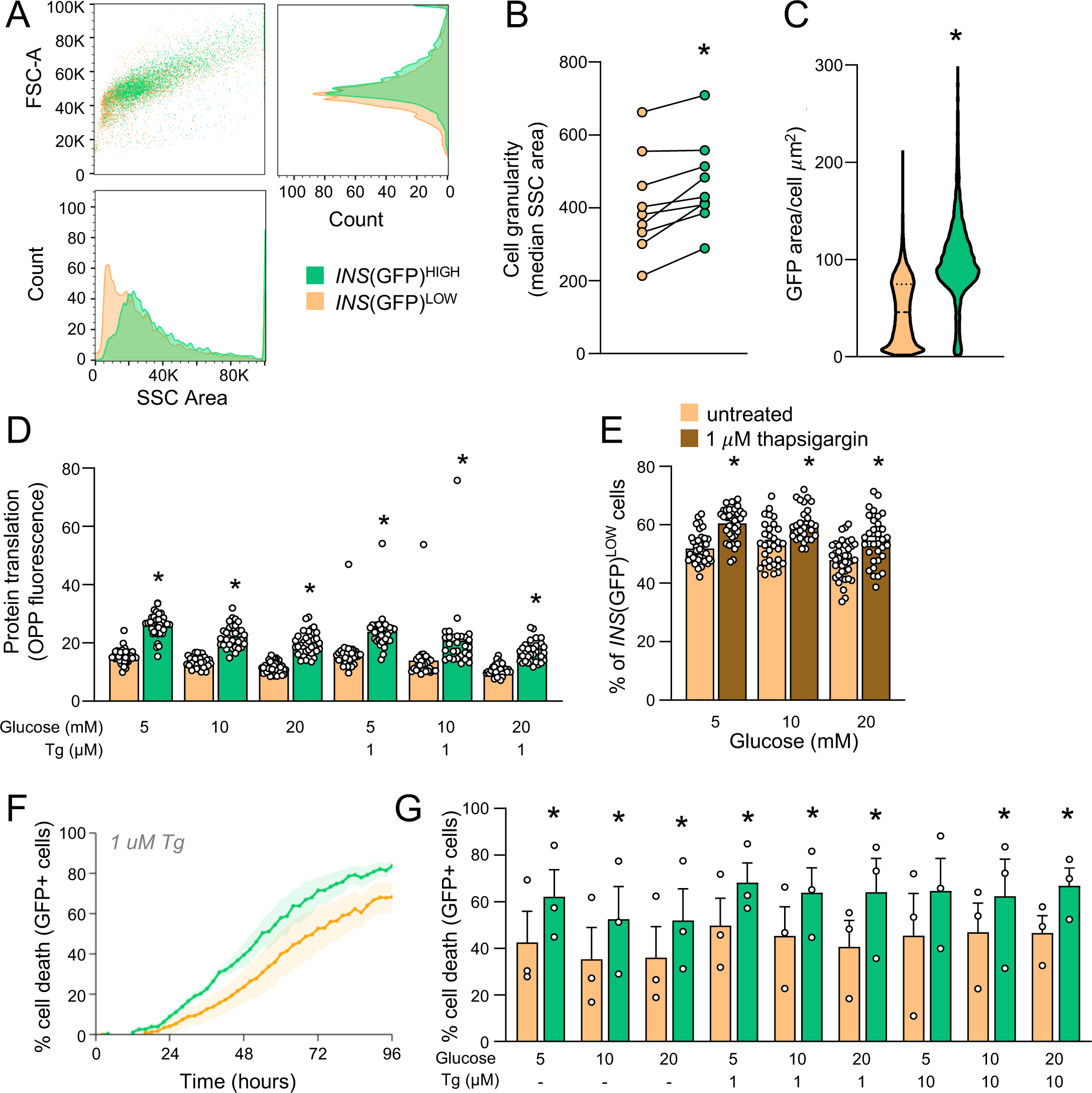
Functional assays of *INS* states in primary β cells. **(A)** FACS data depicting FSC-A and SSC-A of *INS*(GFP)^HIGH^ and *INS*(GFP)^LOW^ primary β cells. **(B)** Quantification of median SSC reflecting granularity. **(C)** Quantification of cell size in plated down cells in the context of *INS*(GFP)^HIGH^ and *INS*(GFP)^LOW^ states using GFP fluorescence area. Student’s t-test. **(D)** OPP protein synthesis assay of *INS*(GFP)^HIGH^ and *INS*(GFP)^LOW^ states in various culture conditions. Data was normalized to cell size. Student’s t-test (n=30-40 wells). **(E)** Ratios of *INS*(GFP)^LOW^ states analyzed in the OPP assay with and without Tg (n=30-40 wells). **(F)** Live Imaging over 96 hours at 30-minute intervals in the indicated conditions (5 mM glucose). Cell death was measured using propidium iodide cell and averaged across replicates. Shading is SEM. (n=human islet 3 donors). **(G)** Quantification of cell death after 72 hours of imaging. Tg: Thapsigargin. Paired t-test. * p < 0.05

### patchSeq, scRNAseq, and SCENIC analysis of primary human β cells in low and low INS states

We leveraged published patchSeq data^28–30^ from 916 β cells across 64 non-diabetic donors to examine the electrophysiological properties of primary β cells in the low and high insulin production states (Fig. 3A). The size of the patchSeq dataset precluded a within-donor high/low classifications, so *INS* mRNA expression was treated as a continuous variable and correlated directly with electrophysiological properties sign-corrected such that higher values reflect greater activity. While effect sizes were small (|r| ≤ 0.1), the associations were directionally coherent. *INS* expression was positively correlated with late Ca^2+^ current amplitude, Ca^2+^ integral, peak Na^+^ current amplitude, and early Ca^2+^ current amplitude, alongside reduced hyperpolarization-activated current, illustrating a subtle shift toward greater membrane excitability and depolarization capacity (Fig. 3A).

**Figure 3.**
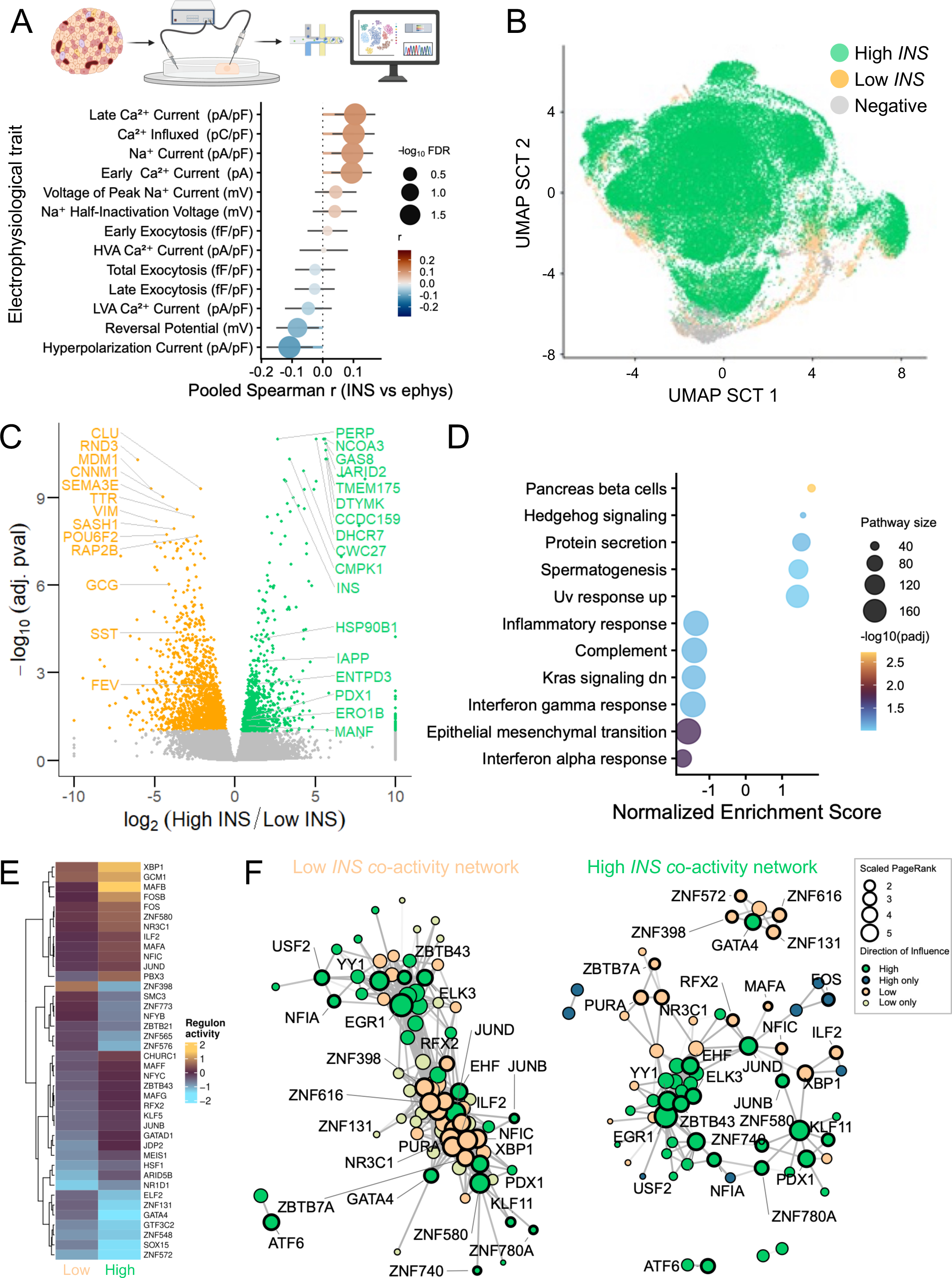
Single cell RNA sequencing analysis of *INS* production states in primary β cells. **(A)** Pooled Spearman correlations between *INS* mRNA expression and electrophysiological properties in patchSeq beta cells (n = 916 cells, 64 donors). Effect sizes represent pooled within-donor correlations using random effects meta-analysis with Hartung-Knapp variance inflation. Points are sized by – log_10_(FDR); colors indicate correlation direction. Electrophysiological variables were sign-corrected such that positive values reflect greater current activity. **(B)** UMAP of cells in the three *INS* states. **(C)** volcano plot of differentially expressed genes comparing high and low *INS* cells. **(D)** Gene set enrichment analysis (GSEA) of Hallmark pathways in high vs low *INS* β cells, using depth-corrected pseudobulk differential expression statistics as the ranking metric. Pathways with p.adj < 0.1 are shown. Point size reflects pathway gene set size; color indicates –log_10_(p.adj). **(E)** Mean SCENIC regulon activity scores for the top 40 most differentially active regulons between high and low *INS* β cells, shown as mean activity per group. Regulons are clustered by similarity. **(F)** Transcription factor co-activity networks for low *INS* (top) and high *INS* (bottom) β cells, derived from bootstrap consensus correlation of SCENIC AUC scores across 100 subsampled iterations. Node size reflects mean PageRank centrality; node color indicates direction of influence relative to the opposing group. Selected transcription factors of interest are labeled.

The transcriptional profiles of low and high *INS* producing cells was examined in detail using a larger dataset comprising 60,663 human β cells from 69 non-diabetic donors (Fig. 3B). We used *INS* mRNA as a surrogate for *INS* gene activity and insulin production, given the high correlation between endogenous *Ins2* (GFP) gene activity and *Ins2* mRNA in our previous scRNAseq analysis of β cells from *Ins2*^GFP^ knock-in mice^20^. We categorized β cells into high, low and negative *INS* mRNA groups within each donor (Fig. 3B). Counts were aggregated into pseudobulk profiles per donor per *INS* expression group to perform differential gene expression analysis. The most significantly upregulated genes in *INS* high β cells were: PERP, a transmembrane protein and a target of p53 in the context of apoptosis^31^; NCOA3, a promoter of cell survival and has been linked to melanoma progression^32^; DHCR7, encoding for the 7-dehydrocholesterol reductase enzyme which catalyzes the final step of cholesterol biosynthesis^33^; GAS8, a cilia motor protein known to be expressed in islet cells^34^; and JARID2, a gene required for the differentiation program of β cells^35^ (Fig. 3C). Cells with high *INS* mRNA had higher expression of genes associated with β cell maturity, including *IAPP*, *ENTPD3* and *PDX1*, as well as coordinated upregulation of multiple ER protein folding and quality control genes, including *ERO1B*, *HSP90B1*, *HSPA5*, and *CANX*, consistent with the high biosynthetic demand of active insulin production (Fig. 3C). Gene set enrichment analysis identified ‘pancreas beta cells’, ‘hedgehog signaling’, and ‘protein secretion’ as the most top positively associated pathways in high *INS* β cells (Fig. 3D). Upregulation of these pathways suggest high *INS* β cells are more functionally mature.

During the analysis, we found that *INS* mRNA level was moderately correlated with total transcript count per cell (Spearman rho = 0.40), and thus we included mean library size as a covariate to ensure that identified differences reflected genuine transcriptional variation rather than sequencing depth (Supplemental Fig. 5A-F). We also examined genes which were positively and negatively correlated to *INS* mRNA using within-donor Spearman rank correlation meta-analysis across studies using a random effects model with Hartung-Knapp variance inflation. *INS* expression was moderately correlated with total transcript count per cell even after excluding cells with no detectable *INS* expression, and inspection of top correlated genes revealed that many showed substantial correlation with library depth, including many ribosomal genes. When not including genes strongly correlated with library depth, top genes correlated with *INS* mRNA include an enhancer of insulin secretion *ADCYAP1*^31^, β cell maturity and functional markers and an ER stress factors (Supplemental Fig. 5). These data demonstrate that cells in the high *INS* state have transcriptomic profiles associated with increased maturity, functionality, but also with ER stress. These results were not significantly affected by donor variables (Supplemental Fig. 6), indicating that they are consistent across age, sex and BMI.

We used SCENIC^26^ to predict transcription factor activity in low and high *INS* β cells. Mean regulon activity scores for the top 40 most differentially active regulons revealed distinct transcriptional signatures between groups (Fig. 3E). Low *INS* cells showed higher activity of zinc finger protein regulons including *GATA4*, *GTF3C2*, *ZNF548*, *SOX15*, and *ZNF572*, while high *INS* β cells showed elevated activity of stress-response and β cell identity regulons including *XBP1*, *MAFA*, *MAFB*, *FOS*, *JUND*, and *NR3C1* (Fig. 3E; Supplemental Fig. 7). Co-activity network topology in low *INS* β cells had higher mean degree (median 28 vs 10), more edges (median 1100 vs 338), and higher density but lower modularity (0.39 vs 0.50; all FDR < 0.001), indicating more uniform connectivity without the discrete hub-and-spoke community structure seen in high *INS* cells (Fig. 3F). High *INS* β cell networks formed tightly organized communities around transcription factors that regulate stress, including a module with *GATA4*, and were characterized by higher modularity and clustering coefficients relative to low INS β cells (Fig. 3F, Supplemental Fig. 7). The presence of distinct node communities and direction-specific hubs reinforces the idea that *INS* expression level stratifies β cells into functionally distinct regulatory states. Overall, our single-cell transcriptomic analyses reveal that high *INS* β cells are enriched for markers of maturity and ER stress and possessed coordinated transcription factor regulatory networks revolving around key stress and β cell transcription factor predicted activities.

### Proteomic profiling of human primary β cells in low and high INS production states

To complement and validated the mRNA data^19^, we performed mass-spec based proteomics on β cells from 13 human donors which were FACS purified into the high, low and negative *INS* states/populations (Fig. 4A). We first took the opportunity to define proteins associated with *INS*(GFP)^HIGH^ β cells compared to *INS*(GFP)^NEG^ non-β cells, and found that β cells are highly enriched in: S-phase cyclin A associated protein in the ER (SCAPER), a ubiquitous protein which plays a role in the cell cycle and is part of a ISL1-SCAPER genomic regulatory block^36^; cyclin dependent kinase inhibitor 1C (CDKN1C), a variant of which has been linked to early-adulthood-onset diabetes and whose demethylation has been shown to induce human β cell replication and^37^; and leucine rich repeat and fibronectin type III domain containing 2 (LRFN2), a protein linked to type 2 diabetes and insulin secretion in β cells^38^ (Fig. 4B). This dataset can be a rich resource for β cell purification targets, using either positive or negative selection strategies.

**Figure 4.**
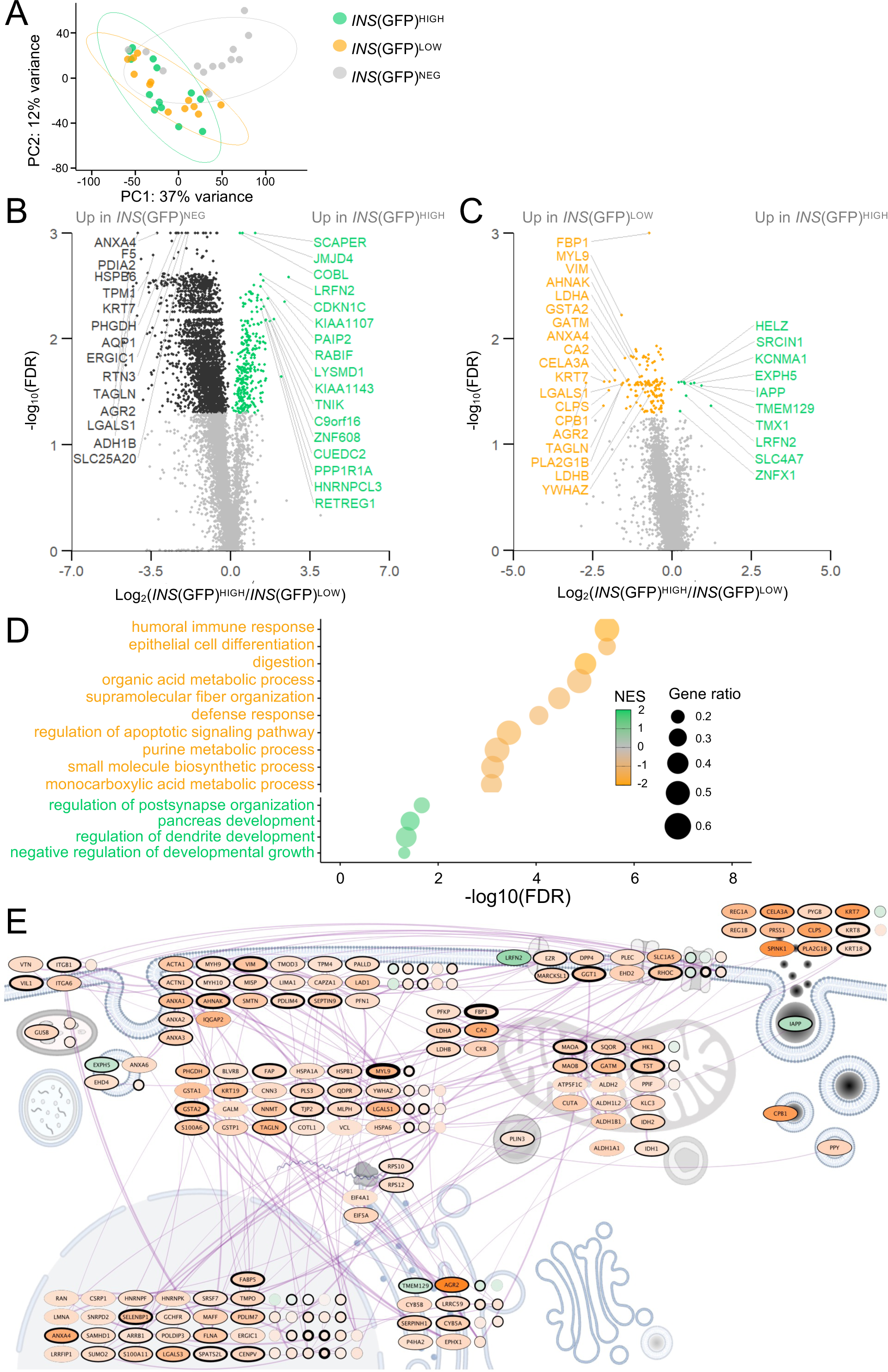
Proteomic analysis of *INS*-GFP primary β cells FACS purified into insulin production states. **(A)** Principal component analysis of protein profiles of FACS purified primary β cells transduced with an *INS*-GFP adenovirus (n=13 donors). **(B)** Volcano plot of differentially expressed proteins comparing FACS purified *INS*(GFP)^HIGH^ and *INS*(GFP)^NEG^ primary β cells. **(C)** Volcano plot of differentially expressed proteins comparing FACS purified *INS*(GFP)^HIGH^ and *INS*(GFP)^LOW^ primary β cells. **(D)** Gene set enrichment analysis of pathways enriched in *INS*(GFP)^HIGH^ and *INS*(GFP)^LOW^ primary β cells. **(E)** Protein-protein interaction network of differentially abundant proteins in the context of cellular compartments. * FDR < 0.05

The top proteins that were more abundant in *INS*(GFP)^HIGH^ β cells compared to *INS*(GFP)^LOW^ β cells included helicase with zinc finger (HELZ), a positive regulator of translation initiation and ribosomal protein S6 phosphorylation^39^; SRC kinase signaling inhibitor 1 (SRCIN1, encodes for the p140Cap protein), a β cell-enriched protein linked to insulin secretion^40^; and potassium calcium-activated channel subfamily M alpha 1 (KCNMA1), a calcium activated BK channel believed to regulate insulin secretion through action potential repolarization (Fig. 4C). The β cell maturity marker islet amyloid polypeptide (IAPP) was strongly upregulated at the protein level in *INS*(GFP)^HIGH^ β cells^41^, as was thioredoxin-related transmembrane protein 1 (TMX1), an oxidoreductase which reversibly oxidizes in response to protein accumulation in the endoplasmic reticulum^42^ (Fig. 4C). Proteins more abundant in *INS*(GFP)^LOW^ β cells relative to *INS*(GFP)^HIGH^ β cells include fructose-bisphosphatase 1 (FBP1), a protein that reduces insulin secretion when overexpressed in β cells^43^; myosin light chain 9 (MYL9), a protein found to be significantly reduced in older mice islets compared to younger^44^; and vimentin (VIM), a protein found to be increased in α and β cells in type 2 diabetes and has been shown to play a role in β cell regeneration^45^. Other proteins that are more abundant in *INS*(GFP)^LOW^ cells include 14-3-3-zeta (YWHAZ), a protein known for constraining β cell insulin secretion by mitochondrial regulation^46^; lactate dehydrogenase A (LDHA), a protein which can induce β cell dedifferentiation in the context of type 2 diabetes^47^; and lactate dehydrogenase B (LDHB), a regulator of basal insulin secretion and lactate levels^48^. Gene set pathway analysis showed that ‘postsynapse organization’, ‘pancreas development’, ‘dendrite development’, and ‘negative regulation of developmental growth’ were enriched in *INS*(GFP)^HIGH^ β cells (Fig. 4D). *INS*(GFP)^LOW^ β cells exhibited enrichment of ‘immune response’, ‘epithelial cell differentiation’, and ‘digestion pathways’. We used STRING and generated protein-protein interaction networks (PPI) to better visualize cellular compartment context. Differentially abundant protein networks were located at the cell surface, nuclei, and mitochondria (Fig. 4E). This deep proteomics dataset identifies novel human β cell markers and further demonstrates that *INS*(GFP)^HIGH^ β cells are enriched with proteins that promote β cell function.

### Functional characterization of SCβ cells in low and high INS states

Similar to primary human β cells, FACS and live cell imaging analysis revealed high *INS* SCβ cells had greater size and granularity compared with low *INS* SCβ cells, (Fig. 5A-C). Normalized to cell size, OPP incorporation showed that Day 35 SCβ cells in the *INS*(EGFP)^HIGH^ state have significantly higher translational activity across multiple culture conditions (Fig. 5D, Supplemental Fig. 8A). Expectedly, global translation was blunted under ER stress conditions, and increased by high glucose and the insulin-transferrin-selenium supplement (Fig. 5D). FACS-purified *INS*(EGFP)^HIGH^ SCβ cells exhibited increased basal and nutrient-stimulated C-peptide release in static incubation (Fig. 5E). Glucagon secretion across these same conditions was not significantly different between the high and low *INS* SCβ cells (Supplemental Fig. 8B). SCβ cell survival was assessed in 48 conditions with the *INS*(EGFP)^HIGH^ state exhibiting consistently higher cell death than *INS*(EGFP)^LOW^ SCβ cells (Fig. 5F-H). These results corroborate our findings in the human primary β cells, revealing that high *INS* SCβ cells are larger, more translationally active, secrete more insulin, but are more fragile in the context of diabetes associated stresses.

**Figure 5.**
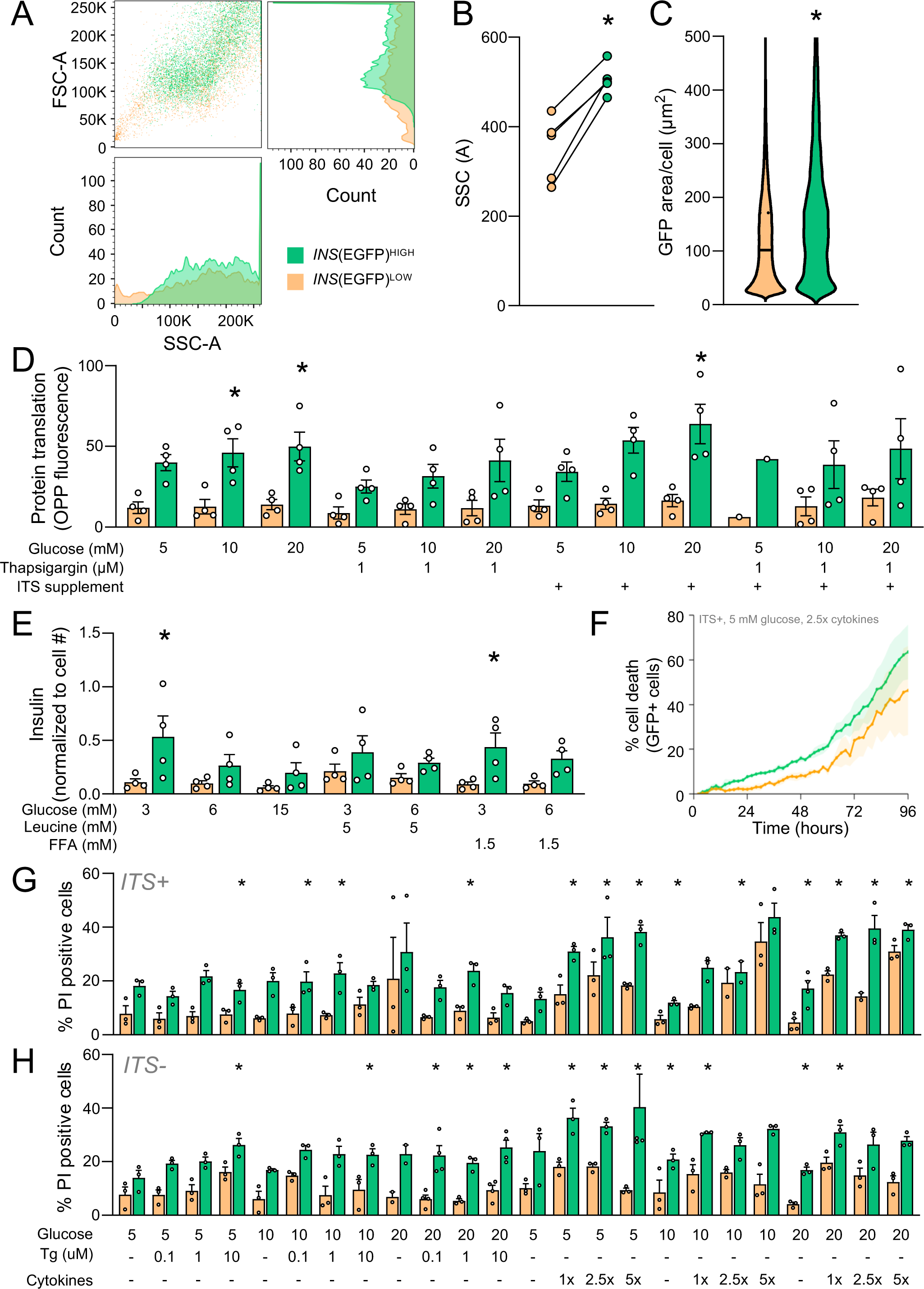
Functional assays of *INS* production states in SCβ cells. **(A)** FACS data depicting FSC-A and SSC-A of *INS*(EGFP)^HIGH^ and *INS*(EGFP)^LOW^ SCβ cells. **(B)** Quantification of median SSC reflecting granularity. **(C)** Quantification of cell size in plated down cells in the context of *INS*(EGFP)^HIGH^ and *INS*(EGFP)^LOW^ states using GFP fluorescence area. **(D)** OPP protein synthesis assay of *INS*(EGFP)^HIGH^ and *INS*(EGFP)^LOW^ states in various culture conditions (n=4 differentiations). Data was normalized to cell size. **(E)** Static incubation insulin secretion measurements of SCβ cells dispersed and FACS purified into *INS*(EGFP)^HIGH^ and *INS*(EGFP)^LOW^ states. ANOVA. **(F)** Live imaging over 96 hours at 30-minute intervals in the indicated conditions (5 mM glucose, with cytokines). Cell death was measured using propidium iodide cell death marker. **(G-H)** Quantification of image-based cell death assay at 72 hours in multiple culture conditions, with and without ITS (n=3 wells). Paired t-test. * p < 0.05

### patchSeq, scRNAseq, and SCENIC analysis of SCβ cells in low and high INS gene activity states

We analyzed patch-seq data from high and low *INS* SCβ cells to link *INS* levels to physiology. *INS*(EGFP)^HIGH^ SCβ cells had reduced Ca^2+^ and Na^+^ charge entry, and lower exocytosis, relatively to *INS*(EGFP)^LOW^ SCβ cells (Fig. 6A-C). On the other hand, transcriptomic comparisons of *INS*(EGFP)^HIGH^ with *INS*(EGFP)^LOW^ cells were generally consistent with our data on primary β cells. *INS*(EGFP)^HIGH^ cells showed upregulation of mRNAs related to β cell maturity and function (*INS*, *PDX1*, *NKX6-1*, *MAFB*, and *PCSK1*), as well as stress markers such as *HSP90B1* (Supplemental Fig. 9A). These data reveal a disconnect between transcription factor and marker mRNA expression and single SCβ electrophysiological maturity. It should be noted that Ca^2+^ current, Na^+^ current, and exocytosis tend to be higher in SCβ compared to primary β cells^25^, so the *INS*(EGFP)^HIGH^ SCβ cells are therefore more functionally similar to primary β cells.

**Figure 6.**
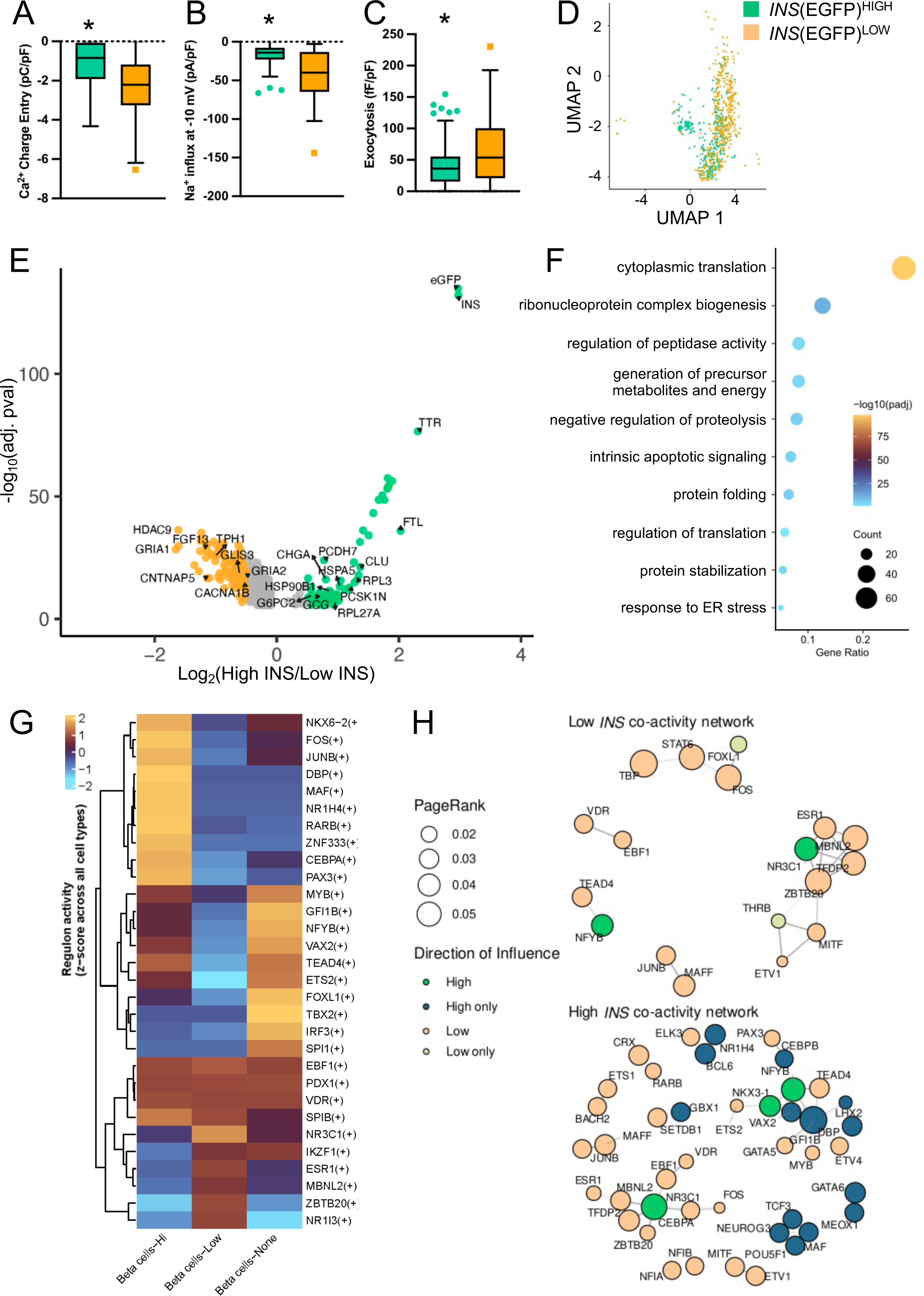
Single cell RNA sequencing analyses of *INS* production states in SCβ cells. **(A-C)** Electrophysiology analyses of *INS*(EGFP)^HIGH^ and *INS*(EGFP)^LOW^ SCβ cells. **(D)** UMAP of *INS*(EGFP)^HIGH^ and *INS*(EGFP)^LOW^ SCβ cells. **(E)** Volcano plot of top differentially expressed genes in *INS*(EGFP)^HIGH^ and *INS*(EGFP)^LOW^ SCβ cells. **(F)** Gene set enrichment analyses comparing *INS*(EGFP)^HIGH^ and *INS*(EGFP)^LOW^ SCβ cells. **(G)** SCENIC analysis for TF activity. Representative heatmap of major TFs governing the various *INS* production states in SCβ cells. **(H)** Transcription factor co-activity networks for *INS*(EGFP)^HIGH^ and *INS*(EGFP)^LOW^ SCβ cells, derived from bootstrap consensus correlation of SCENIC AUC scores across 100 subsampled iterations. Node size reflects mean PageRank centrality; node color indicates direction of influence relative to the opposing group. Selected transcription factors of interest are labeled.

To further understand the molecular mechanisms underlying functional differences between SCβ cells in the low and high insulin production states we further analyzed scRNAseq data. Like EGFP fluorescence, Egfp mRNA had a bimodal distribution (Supplemental Fig. 9B). As expected, *INS* mRNA was upregulated in SCβ cells with higher *Egfp* mRNA (Fig. 6E, Supplemental Fig. 9C). Alongside *INS*, the top two other genes were the positive regulator of β cell stimulus secretion coupling transthyretin (*TTR*)^49^, and the iron ferroptosis and oxidative stress regulator ferritin light chain (*FTL*)^50^. We also found enrichment in genes encoding for the ribosomal proteins *RPL3* and *RPL27A*, and the ER chaperone protein heat shock protein family A member 5 (*HSPA5*, encoding for the BIP protein) (Fig 6E, Supplemental Fig. 9C). Genes upregulated in the *INS*(EGFP)^LOW^ cell state include *FGF13*, from the fibroblast growth factor family we have recently shown are highly effective at protecting SCβ cells^51^, as well as the histone protein *HDAC9*, and the glutamate inotropic receptor AMPA type subunit 1 (*GRIA1*) (Fig. 6E, Supplemental Fig. 9C). Gene set enrichment analysis revealed that the *INS*(EGFP)^HIGH^ state was associated with ‘cytoplasmic translation’, ‘ribonucleoprotein complex biogenesis’, and ‘regulation of peptidase activity’ (Fig. 6F). Also consistent with function assays (Fig. 5), ‘protein folding’ and ‘response to ER stress’ were upregulated (Fig. 6F).

As a complementary analysis, we asked which genes had the highest positive correlation with *Egfp* mRNA, analyzed as a continuous variable (Supplemental Fig. 10). As expected, *INS* had the highest correlation to *Egfp*, followed by *TTR* (Supplemental Fig. 10). Other correlated genes of interest included the aforementioned *Hspa5*; another ER protein folding gene heat shock protein 90 β family member 1 (*HSP90B1*); the β cell maturity marker chromogranin A (*CHGA*); the insulin secretion regulator glucose-6-phosphatase catalytic subunit 2 (*G6PC6*)^52^; and protocadherin 7 (*PCDH7*), a gene shown to select for SCβ cells with enhanced insulin secretion^53^ (Supplemental Fig. 10). Ribosome components and mitochondrial genes were also positively correlated (Supplemental Fig. 10). Genes negatively correlated with *Egfp* mRNA included the *INS* gene pioneering factor GLIS family zinc finger 3 (*Glis3*)^54^, a gene that is associated with both type 1 diabetes and type 2 diabetes risk, and is known to play a protective role in β cells^55^ (Supplemental Fig. 11). *Egfp* mRNA was also negatively correlated with expression of the N-type calcium voltage-gated channel subunit alpha1 B (*CACNA1B*); glutamate receptors *Gria1* and *Gria2*; and neural cell adhesion molecule (*NCAM1*), a protein whose polysialylated form has been shown to be a marker for β cell functionality^56^ (Supplemental Fig. 11).

We also performed differential gene analysis on cells with high and low *INS* mRNA to directly match the human primary β cell analysis and benchmark against the EGFP marker (Supplemental Fig. 12A, B). Consistent with our findings with the *EGFP* analysis, the top genes upregulated in high *INS* SCβ cells were *Egfp*, *TTR*, and *FTL* (Supplemental Fig. 12A). Several of the top genes differentially expressed in the *Egfp* analysis were also found in this analysis, including *RPL3*, *HDAC9*, and *GRIA1* (Supplemental Fig. 12A). Genes set enrichment analysis identified ‘cytoplasmic translation’, ‘protein folding in endoplasmic reticulum’, and ‘hormone transport’ as the top three pathways upregulated in high *INS* SCβ cells (Supplemental Fig. 12B), showing strong consistency with pathways identified in the *Egfp* analysis (Fig. 6F). Together, these scRNAseq analyses concur with the functional assays, showing upregulation insulin production, protein translation, and ER stress response in high *INS* SCβ cells.

To identify transcription factors associated with high and low *INS* SCβ cells, we next performed SCENIC analysis on our scRNAseq data. Similar to findings in the primary human β cell analysis, top transcription factors active in high *INS* SCβ cells include maturity markers such as *NKX6-2* and *PDX1*, as well as stress regulators *FOS* and *JUNB* (Fig. 6G). We also identify transcription factors which were highly active in other cell types in SC-islets, such as the maturity-onset diabetes of the young 5 associated and regulator of pancreatic development HNF1B^54,55^, the developmental regulator of photoreceptor cells *CRX*^56^, and the pancreatic cancer associated *FOSL1*^57^ in enterochromaffin cells; the immune response transcription *STAT1*, the activator of the proglucagon promoter *FOXA3*^58^, and the differentiation master regulator *NEUROG3* in α cells; a regulator of α cell identity regulation *KLF4*^59^, and the type 2 diabetes driver *BACH2* in SST-positive cells^60^ (Fig. 6G). Network analysis revealed *INS*-state-associated differences in transcription factor co-activity within SCβ cells, though the nature and magnitude of these differences differed substantially from primary β cells (Fig. 6H, Supplemental Fig. 13). Direct topological comparison between the two datasets is methodologically limited by independent SCENIC runs, different co-activity thresholds (ρ ≥ 0.45 for SCβ vs ρ ≥ 0.6 for primary β cells), and fewer detected regulons in SCβ cells (127 vs 313), including the absence of key maturity-associated regulons such as *JUND*, *MAFA*, *MAFB*, and *SREBF1/2* (Fig. 6H). Within SCβ cells, high *INS* networks had significantly more nodes and edges than low *INS* networks (median 41 vs 32 nodes, 49 vs 38 edges; both FDR < 0.001), with higher degree skew and modularity, consistent with a more selectively organized network structure (Supplemental Fig. 13A). In contrast, low *INS* SCβ networks were denser with higher clustering coefficients (FDR < 0.001), indicating tighter local connectivity among a smaller set of co-active regulons (Fig. 6H). Mean degree did not differ significantly between groups (p = 0.74), suggesting that individual transcription factors maintain similar connectivity on average despite differences in overall network size and organization (Supplemental Fig. 13B). These more subtle differences in network topology between *INS* states in SCβ cells, compared to the extensive rewiring observed in primary β cells, suggest that *INS* production-associated transcriptional network organization may not be fully established in current SCβ differentiation protocols.

### Proteomic analysis of INS states in SCβ cells

To further characterize high *INS* SCβ cells, we performed mass-spec based proteomics on FACS purified SCβ cells differentiated for 35 and 50 days with protocols A and B^18,57^. Single-cell proteomics is not yet feasible at the scale of scRNAseq, so we sorted cells into high EGFP, low EGFP, and negative EGFP populations (Fig. 7A). Interestingly, PCA analyses revealed that day 35 high *INS* SCβ cells were more similar to day 50 low *INS* SCβ cells (Fig 7A). We compared protein abundance between *INS*(EGFP)^HIGH^ with *INS*(EGFP)^NEG^ cells (Supplemental Fig. 14A-C). In S7 day 35 cells, proteins enriched in *INS*(EGFP)^HIGH^ included the β cell marker ENTPD3, the stress marker ERO1B, and a marker for islet precursor cells DMBT1^58^. In S7 day 50 cells, top proteins in high *INS* SCβ included ENTPD3, β cell function marker PCDH7^59^, and kynurenine 3-monoxygenase (KMO), a protein known to be upregulated under metabolic stress in islets^60^. Across both S7 day 35 and S7 day 50 cells, we observed many well-known ER proteins (HSPA5, HSP90B1, MANF) that were upregulated high *INS* SCβ cells, as well as β cell maturity markers IAPP and PDX1 (Supplemental Fig. 14A, B). A common downregulated protein in high *INS* SCβ cells of both ages was annexin A2 (ANXA2), a protein which binds the insulin receptor^61^ and plays a role in cell growth (Supplemental Fig. 14A, B). Gene set enrichment analysis revealed that S7 day 50 high *INS* SCβ had upregulation of ‘rRNA processing’, ‘oxidative phosphorylation’, and ‘mitochondrial translation’ (Supplemental Fig. 14C). This suggests that high *INS* SCβ cells differ from other cell types in SC-islets primarily in pathways related to mitochondria activity and oxidative respiration. High *INS* SCβ cells exhibited upregulation of ‘ER to Golgi vesicle mediated transport’ and ‘ERAD pathway” (Supplemental Fig. 14C).

**Figure 7.**
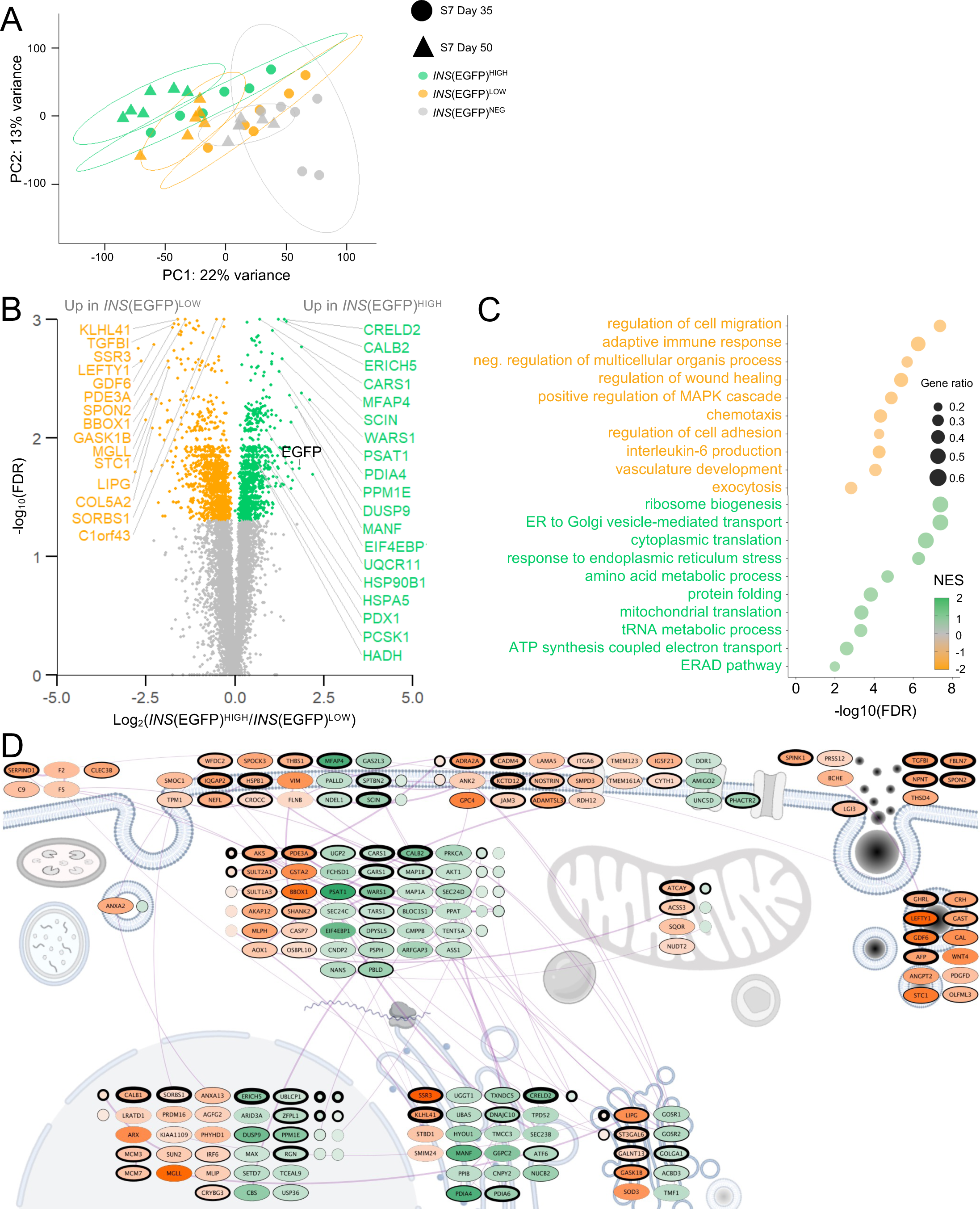
Proteomic analysis of *INS*-EGFP SCβ cells FACS purified into insulin production states cultured using protocol B. **(A)** Principal component analysis of protein profiles of FACS purified S7 day 35 and S7 day 50 *INS*-EGFP SCβ cells (n=6 differentiations per age). **(B)** Volcano plot of differentially expressed proteins comparing FACS purified S7 day 50 *INS*(EGFP)^HIGH^ and *INS*(EGFP)^low^ SCβ cells. **(C)** Gene set enrichment analysis of pathways enriched in S7 day 50 *INS*(EGFP)^HIGH^ and *INS*(EGFP)^LOW^ SCβ cells. **(D)** Protein-protein interaction network of differentially abundant proteins in the context of cellular compartments * FDR < 0.05

Comparison of *INS*(EGFP)^HIGH^ with *INS*(EGFP)^LOW^ SCβ cells cultured with protocol B revealed that maturity markers PDX1 and ENTPD3 were upregulated at both day 35 and day 50 (Fig. 7B, Supplemental Fig. 15), as well as the insulin processing protein PCSK1. We observed upregulation of multiple ER proteins (HSPA5, HSP90B1, MANF) in high *INS* SCβ cells at both ages (Fig. 7B, Supplemental Fig. 15). The protein disulfide isomerase family A member 4 (PDIA4), which plays roles in β cell dysfunction and death in diabetes, was upregulated in both S7 day 35 and S7 day 50 high *INS* SCβ cells^62^ (Fig. 7B, Supplemental Fig. 15). Gene set enrichment revealed that pathways upregulated in high *INS* SCβ cells included ‘ribosome biogenesis’, ‘ER to Golgi vesicle-mediated transport’, and ‘cytoplasmic translation’ (Fig. 7C). Importantly, ‘ER to Golgi vesicle-mediated transport’ and ‘ERAD’ pathway were also major pathways upregulated in high *INS* SCβ cells when comparing *INS*(EGFP)^HIGH^ with *INS*(EGFP)^NEG^ (Fig. 7C, Supplemental Fig. 14C).

We profiled the proteomes of FACS purified SCβ cells cultured for 35 or 50 days using protocol A. Major ER proteins (HSPA5, HSP90B1, MANF, PDIA4) were again upregulated in high *INS* SCβ cells of both ages relative to *INS*(EGFP)^NEG^ cells (Supplemental Fig. 16A, B). The mitochondrial pyrroline-5-carboxylate reductase 1 (PYCR1) was also upregulated in high *INS* SCβ cells (Supplemental Fig. 16A, B). Common downregulated proteins in high *INS* included ANXA2 and members of the glutathione S-transferase family (GSTA), which play roles in detoxification of electrophilic compounds (Supplemental Fig. 16A, B). Gene set enrichment analysis revealed ‘Golgi vesicle transport’, ‘ER to Golgi vesicle-mediated transport’, and ‘mitochondrial translation’ as top pathways upregulated in high *INS* SCβ cells, complementing parallel analysis of protocol B cells (Supplemental Fig. 16C).

Comparison of *INS*(EGFP)^HIGH^ with *INS*(EGFP)^LOW^ SCβ cells cultured for 35 days or 50 days using protocol A revealed similar results to protocol B (Supplemental Fig. 17A, B). ER proteins (HSPA5, HSP90B1, MANF) were again a major group of proteins with greater abundance in high *INS* SCβ cells of both ages, as were β cell maturity markers PDX1, ENTPD3, and IAPP (Supplemental Fig. 17A, B). Crystallin β A2 (CRYBA2), a β cell immaturity marker that was shown to be upregulated in response to FOXO1 inhibition, was reduced in high *INS* SCβ cells of both ages^63^; as was SLC18A1, an enterochromaffin cell marker (Supplemental Fig. 17A, B). Gene set enrichment analysis revealed ‘ER to Golgi vesicle-mediated transport’, ‘response to endoplasmic reticulum stress’, and ‘protein folding’ as top pathways in S7 day 50 high *INS* SCβ cells, similar to results in our scRNAseq data and protocol B proteomics (Supplemental Fig 17C).

PPIs and cellular compartment analysis illustrated that networks of differentially abundant proteins in high *INS* SCβ cells were common in cytoplasm, nuclei, and ER (Fig. 7D). Overall, across all comparisons, we find that high *INS* SCβ cells had upregulation for major ER proteins and β cell maturity markers, while negative or low *INS* cells had markers of β cell immaturity. ER, Golgi, mitochondria, protein production and stress management pathways were consistently upregulated in high *INS* SCβ cell. These results provide protein-level explanations for functional differences. These findings support the idea that high *INS* SCβ cells are more mature in some ways, but also more fragile.

### Proteomic comparison of high INS primary and SCβ cells

Previous proteomic analysis comparing human islets to stem cell derived islet-like clusters is confounded by significant differences in cell composition^19^. To better identify gaps between primary and stem cell derived β cells that could guide improvements to differentiation protocols, we compared the proteomes of purified primary human high *INS* β cells against purified high *INS* protocol B SCβ cells (Fig. 8A-F). High *INS* SCβ cells were significantly larger than high *INS* primary β cells (Fig. 8A). Principal component analysis of proteomic data illustrated that primary β cells had higher variability, associated with known donor-donor heterogeneity^19,64^ (Fig. 8B). Hierarchical clustering revealed striking overall differences between primary β cells and SCβ cells (Fig. 8C). Nevertheless, key marker proteins generally were directionally consistent between high and low INS cells in both primary and stem cell derived β cells (Supplemental Fig. 18). There was significantly more INS protein in primary β cells than SCβ cells (Fig. 8D).

**Figure 8.**
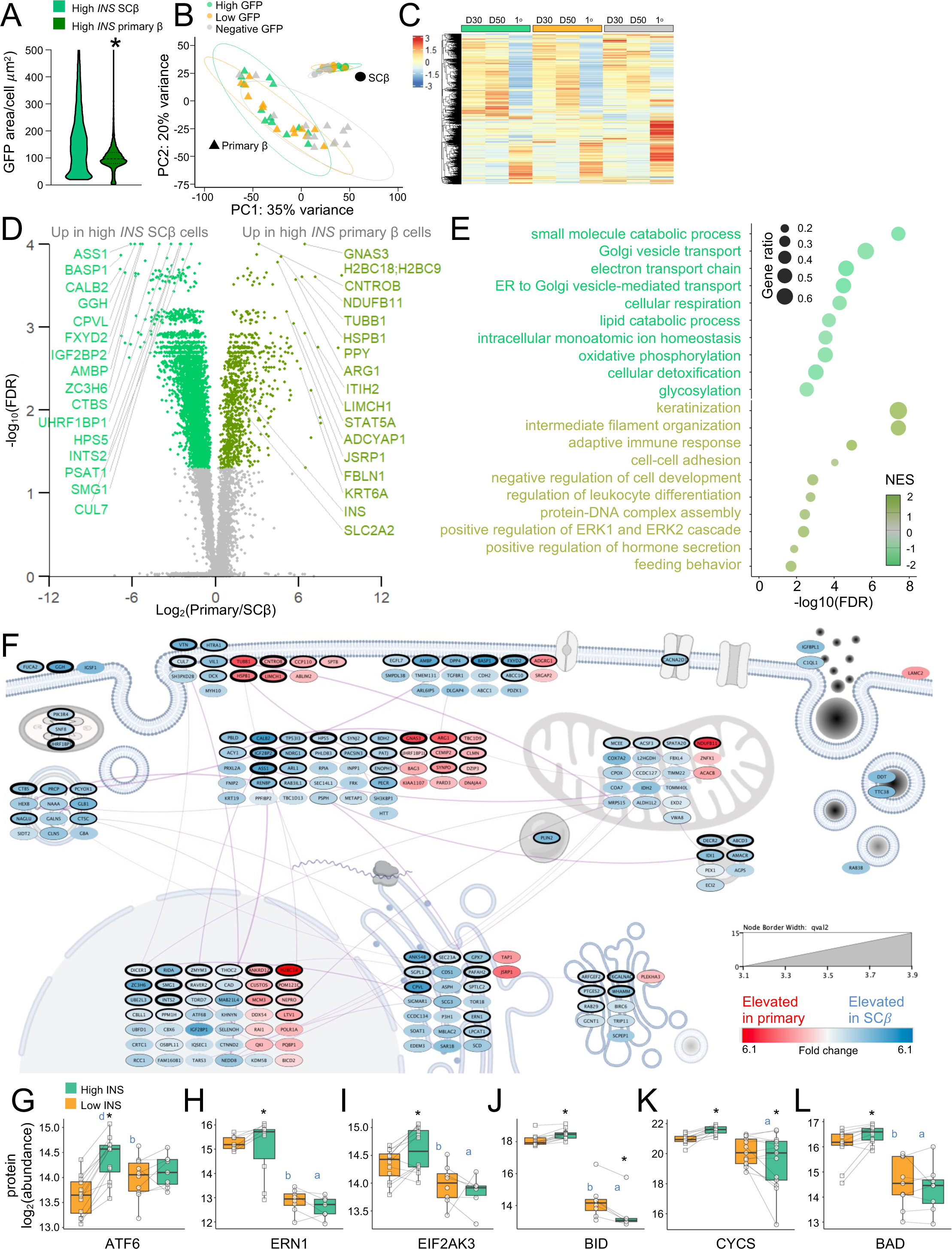
Proteomic comparison of high *INS* states in SCβ cells and human primary β cells. **(A)** Quantification of cell size in plated down cells in the context of SCβ cells and human primary β cells in the high *INS* state using GFP fluorescence area. **(B)** Principal component analysis of protein profiles of FACS purified cells from S7 day 50 *INS*-EGFP SCβ cells (protocol B) and *INS-*GFP human primary β cells (n=6 differentiations, n=13 donors). **(C)** Hierarchical clustering of SCβ cells cultured using protocol B and primary β cells. **(D)** Volcano plot of differentially expressed proteins comparing SCβ cells and primary β cells in the high *INS* state. **(E)** Gene set enrichment analysis of pathways enriched in SCβ cells and primary β cells in the high *INS* state. **(F)** Protein-protein interaction network of differentially abundant proteins in the context of cellular compartments * FDR < 0.05**. (G-L)** Abundances of key UPR and cell death marker proteins from purified day 50 SCβ cells (protocol A and B) and purified primary β cells. * Multiple paired t-test q < 0.05. “a” 2-way ANOVA post-hoc difference between primary β and SCβ (high INS). “b” 2-way ANOVA post-hoc difference between primary β and SCβ (low INS). “c” 2-way ANOVA post-hoc difference between high and low INS (primary β). “d” 2-way ANOVA post-hoc difference between high and low INS (SCβ).

The top proteins upregulated in the primary β cells were GNAS3, an important protein which regulates insulin secretion capacity in β cells^65^; centrobin (CNTROB), a protein important for maintaining centrosomal function known to be highly expressed in β cells; and heat shock protein β-1 (HSPB1), a protein shown to be protective of β cells in the context of proteotoxic and ER stress^66^ (Fig. 8D). Gene set enrichment analysis revealed that high *INS* primary β cells had upregulation of pathways including ‘keratinization’, ‘intermediate filament organization’, and ‘adaptive immune response’ (Fig. 8E). Interestingly, ‘positive regulation of ERK1 and ERK2 cascade’ was also upregulated, possibly suggesting increased insulin signaling as a result of the higher abundance of INS protein (Fig. 8E); prior studies have shown that reducing insulin production downregulates ERK1 and ERK2 in mice^67^. Upregulation of the pathways ‘positive regulation of hormone secretion’ and ‘feeding behavior’ suggest that primary high *INS* β cells differed from their SCβ cell counterparts through secretion mechanisms, consistent with the upregulation of proteins such as GNAS; adenylate cyclase activating polypeptide 1 (ADCYAP1, encodes for PACAP)^68^; and solute carrier family 2 member 2 (SLC2A2), the GLUT2 transporter (Fig. 8E).

Top proteins upregulated in the high INS SCβ cells relative to primary high INS β cells were gamma-glutamyl hydrolase (GGH) a regulator of folate metabolism^69^; argininosuccinate synthase 1 (ASS1) an enzyme important for arginine metabolism in β cells^70^; and FXYD domain containing ion transport regulator 2 (FXYD2), a protein shown to be a marker for β cell function^71^ (Fig. 8D). Gene set enrichment revealed ‘small molecule catabolic process’, ‘Golgi vesicle transport’, and ‘electron transport chain’ as top pathways upregulated in high *INS* SCβ cells (Fig. 8E). ‘ER to Golgi vesicle-mediated transport’ was also upregulated, which may suggest higher protein production in SCβ cells (Fig. 8E). Indeed, the larger cell size of the SCβ cells supports this hypothesis, as cell size often tracks well with protein synthesis^72^(Fig. 8A, Fig. 2D, Fig. 5D). PPI analyses revealed that proteins upregulated in high INS SCβ cells were prominent in the ER and Golgi (Fig. 8F).

The differential translational capacity between the high *INS* primary β cells and high *INS* SCβ cells, prompted us to query our quantitative proteomics dataset for differences in ER stress management proteins focusing on the three arms of the unfolded protein response (UPR). Indeed, ATF6 was positively associated with high INS production states in SCβ cells and trended towards increased in high INS primary β cells (Fig. 8G). Interestingly, IRE1A (ERN1) and PERK (EIF2AK3) were increased in high INS SCβ cells but trended towards reduced in high INS primary β cells (Fig. 8H,I). Comparing across cell types, SCβ cells had more comparable or even reduced levels of the ATF6 when comparing with primary β cells, but had significantly more IRE1A and PERK (Fig 8H-I). We also queried programmed cell death effectors. The protective factor MCL1 was trending towards increased in high INS cells compared with low INS both cell types (Supplemental Fig. 19). Interestingly, the protective diabetes risk factor GLIS3 was trending towards increased in high INS primary β cells, but trended towards reduced in high INS SCβ cells, correlating with our mRNA data (Supplemental Fig. 19). On the other hand when comparing within cell types, other cell death-promoting proteins BID, BAD, CYCS, CASP9, and STAT2 were significantly increased in high INS SCβ cells but significantly reduced or trended towards reduced in high INS primary β cells (Fig. 8J-L, Supplemental Fig. 19). Comparing across cell types, most of these factors were increased or trending towards increased in SCβ cells compared to primary β cells (Fig. 8J-L). This suggests that SCβ cells manage ER stress differently compared to primary β cells. Overall, our proteomics comparisons of high *INS* primary β cells and high *INS* SCβ cells reveal key differences in mechanisms related to protein production and fragility.

## Discussion

This goal of this study was to compare β cells with low versus high *INS* gene activity from human donors or stem cells. Comprehensive examination of low and high insulin production states at the transcriptomic, proteomic, and functional levels, revealed cellular traits and molecular mechanisms associated with high insulin production including significantly increased fragility. We also provide a side-by-side comparison of primary β cells and SCβ cells in the high *INS* state.

The study of β-cell heterogeneity has been enabled by advances in single-cell transcriptomics, complementing the characterization of morphological and functional heterogeneity^64,73^. Using β cells from *Ins2*^GFP^ knock-in mice, we previously identified two distinct *Ins2* gene activity states and showed that cells could transition between these states over time^13,24^. Here, we show similar *INS* activity states exist in primary human β cells and human SCβ cells. Human β cells in the high *INS* state are larger, produce more protein, and secrete more insulin than those in the low *INS* state. It should be noted that all β cell models used in this study are generally post-mitotic, with human donors ranging between 33 and 66 years of age^74^, and SCβ cells having greatly reduced proliferation in stage 7^18^.

Our data confirm a positive relationship between insulin production and β cell vulnerability that is consistent with previous research^13,15,24^. This likely has translational relevance, given that the L-type calcium channel inhibitor verapamil can protect β cells *in vitro*^75^ and delay type 1 diabetes in clinical trials^75^. Similarly, the Na^+^ channel inhibitor carbamazepine protects β cells from multiple stressors and prevents diabetes in NOD mice^76,77^. Our patch-seq analysis shows that the high *INS* state may be associated with hyper-excitability of primary human β cells, and a more mature electrical phenotype in SCβ-cells. Increased production, modification, and/or presentation of neoautoantigens could contribute to the pathogenesis of type 1 diabetes^2^. Excess insulin production, perhaps driven in part by glucose dependent transitions from the low to the high INS state, may also contribute to the basal hyperinsulinemia that increases the risk of type 2 diabetes^6^. Further studies in the context of diabetic models are needed to confirm these hypotheses.

For diabetes cell therapy, SCβ cells must have the ability to produce large amounts of insulin to meet the demands of a recipient with limited endogenous β cells, but the weight of the evidence suggests that these cells are not yet equivalent to primary human β cells. Here, we directly compared primary human β cells and SCβ cells in their high *INS* states. Beyond the similarities discussed above, this head-to-head analysis revealed many substantial differences. SCβ cells were larger than primary human β cells but had less insulin content. *INS* state dynamics were different, with more cells transitioning from the high *INS* state to the low *INS* state in primary β cells^24^. SCENIC analyses revealed more well connected and distinct transcription factor networks in the high *INS* primary β cells than the high *INS* SCβ cells, suggesting that certain transcription factor synergies have yet to be fully formed in SCβ cells. Collectively, this suggests that primary β cells have better control over insulin production and better adaptability.

Proteomic comparisons between the two high *INS* states in primary and stem cell-derived β cells revealed key differences in protein secretion machinery and protein production between the two systems. GNAS, ADCYAP1, and SLC2A2 were upregulated in high *INS* primary β cells but not in SCβ cells, reflecting key differences in the machinery controlling glucose-stimulated insulin release and underlying inadequacies with current differentiation protocols^19^. ER and Golgi proteins were upregulated in high *INS* SCβ cells, suggesting increased protein synthesis, but INS remained low. Upregulation serine regulators PSAT1 and PHGDH in the high *INS* SCβ cells may reflect production of non-canonical proteins in high *INS* SCβ cells^78^. SCβ cells also appeared to be more active at low glucose. Interestingly, hydroxyacyl-CoA dehydrogenase (HADH, also known as SCHAD), whose deficiency has been linked to hyperinsulinemia in children^79^, was consistently upregulated in high *INS* SCβ cells. Overall, our functional assays and omics datasets cells reveal clear differences between the high *INS* states in primary human β cells and SCβ cells and may suggest different trade-offs to deal with the pressure of high insulin production.

Insulin production is taxing to β cells and drives ER stress even under baseline conditions^15^. Failure to resolve ER stress results in cell dysfunction and death. Our data suggest that differences exist in how primary human β cells and SCβ cells manage ER stress. We propose that β cells can transition between high and low insulin production states as a method of reducing stress, and indeed cell state transition behaviors between primary human β cells and SCβ cells were distinct. SCβ cells transition much less frequently from the high INS to the low INS state. Induction of ER stress with Tg increased high to low INS transitions in primary β cells, but not SCβ cells. Of the three main UPR proteins, only ATF6 was positively correlated with INS production across both cell types. ATF6 is important for stress management and adaptation functions, such as promoting ER expansion^80^. However, the other UPR factors (IRE1A, PERK) and multiple cell death factors (BAD, BID, CYCS, CASP9, and STAT2) had opposite patterns between SCβ cells and primary β cells. Furthermore, SCβ cells had considerably higher levels of these same proteins when compared to primary β cells. Taken together, this may suggest that SCβ cells are not as capable of managing stress. Indeed, our cell death assays show that the cell death ratio of the high and low INS states was much smaller in primary β cells compared with SCβ cells under the same conditions. ‘Intrinsic apoptotic signaling’ was also one of the top pathways enriched in high INS SCβs compared to low INS at the mRNA level. Overall, primary β cells seem to be more efficient at managing stress, while SCβ cells are driven towards apoptosis.

### Limitations and Future Directions

Limitations of our study include the use of adenovirus and a short rat insulin 2 reporter to track insulin production states in primary human β cells. This exogenous construct may impact cellular behaviours and may not fully reflect endogenous insulin production. A caveat of the proteomic data on the sorted cells in the low INS group, is that we cannot exclude that some differentially abundant protein signals are related contamination with highly abundant proteins in non-beta cells. FACS purification is also imperfect. For instance, glucagon was detected in both cell states in both cell models. To provide greater purity, additional markers such as CD49a (ITGA1) and ENTPD3 could be used in future studies^81,82^. In our hands, we found that the GFP positive SCβ cells were strongly enriched for CD49a. Similarly, ENTPD3 mRNA and protein were both consistent elevated in high *INS* states. These orthogonal data provide support to our sorting approach. Similarly, signals in single cell RNA sequencing can result from contamination of ‘individual’ cells with highly expressed RNAs that become ambient early in the workflow. Nevertheless, the use of multiple orthogonal technologies to assess the low and high insulin production states, including long-term live cell imaging of unequivocally individual β cells, gives us added confidence in the overall phenotypes. Despite the limitations, our study provided a comprehensive characterization of low and high *INS* states in human primary and stem cell-derived insulin-producing cells at the transcriptomic, proteomic, and functional level. We identified clear differences between the states, highlighted key differences between the two human β cell models, and identified phenotypes which may play a role in diabetes pathophysiology. Future studies should examine these cell states in the context of disease models.

## Author contributions

CMJC designed studies, performed experiments, analyzed/interpreted data, wrote the manuscript. MEO analyzed and interpreted data, wrote parts of the manuscript. JM performed experiments, analyzed/interpreted data. HC analyzed and interpreted data. AW performed experiments. SYC performed experiments. LTHH produced and shared original multi-omics data. RM designed, supported, and analyzed proteomics experiments. JR designed and interpreted proteomics data. BS performed experiments and analyzed data. NS performed experiments. SM performed experiments. CEE analyzed/interpreted data, wrote parts of the manuscript. WWW supervised work. PEM supervised work. FCL performed experiments, supervised work. JDJ conceived the project, designed studies, interpreted data, edited the manuscript, and is the guarantor of this work.

## Acknowledgments

We thank many colleagues for helpful discussions. This manuscript used data acquired from the Human Pancreas Analysis Program (HPAP-RRID:SCR_016202) Database (https://hpap.pmacs.upenn.edu), a Human Islet Research Network (RRID:SCR_014393) consortium (UC4-DK-112217, U01-DK-123594, UC4-DK-112232, and U01-DK-123716). We thank the organ donors and their families. Human islets for research were provided by the Alberta Diabetes Institute IsletCore at the University of Alberta in Edmonton (www.isletcore.ca) with the assistance of the Human Organ Procurement and Exchange (HOPE) program, Trillium Gift of Life Network (TGLN), BC Transplant, Quebec Transplant and other Canadian organ procurement organizations. Islet isolation was approved by the Human Research Ethics Board at the University of Alberta (Pro00013094). All donors’ families gave informed consent for the use of pancreatic tissue in research. We would also like to thank Dr. Robert Baker and Dr. Tim Kieffer for advice, discussion, and the *INS*-GFP adenovirus.

