## Supplemental figures and tables for "Functional, transcriptomic, and proteomic profiles of human primary and stem cell-derived beta cells in a state of high insulin production and increased fragility"

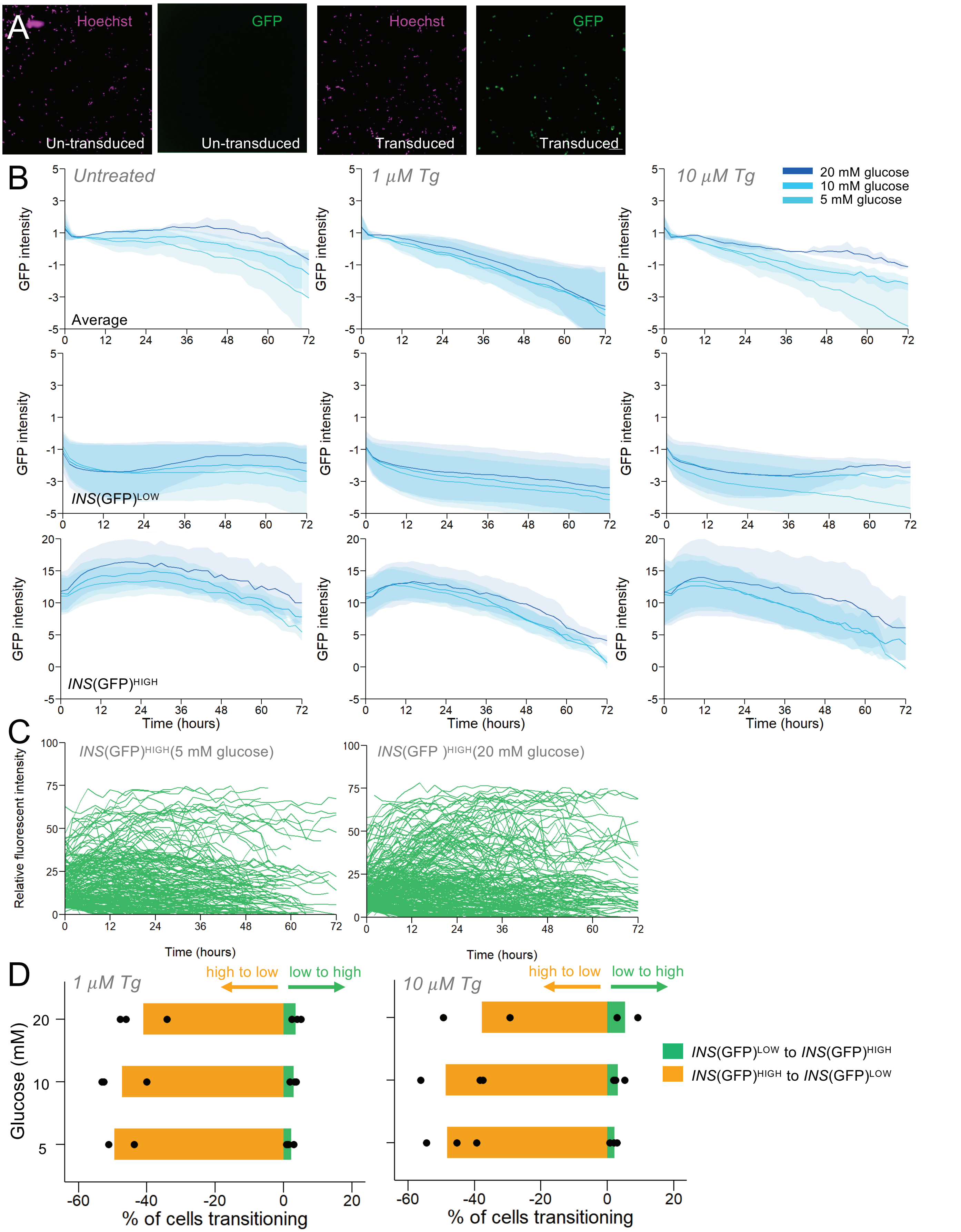


**Supplemental Figure 1. Dynamic *INS* production in human primary β cells. (A)** Representative images of β cells transduced with *INS*-GFP adenovirus. **(B)** GFP fluorescence over time, including average, just *INS*(GFP)$^{LOW}$ and just *INS*(GFP)$^{HIGH}$. **(C)** Traces of cells which started experiments in the *INS*(GFP)$^{HIGH}$ state. Each line = 1 cell. **(D)** Quantification of cell state transitions in the context of thapsigargin.

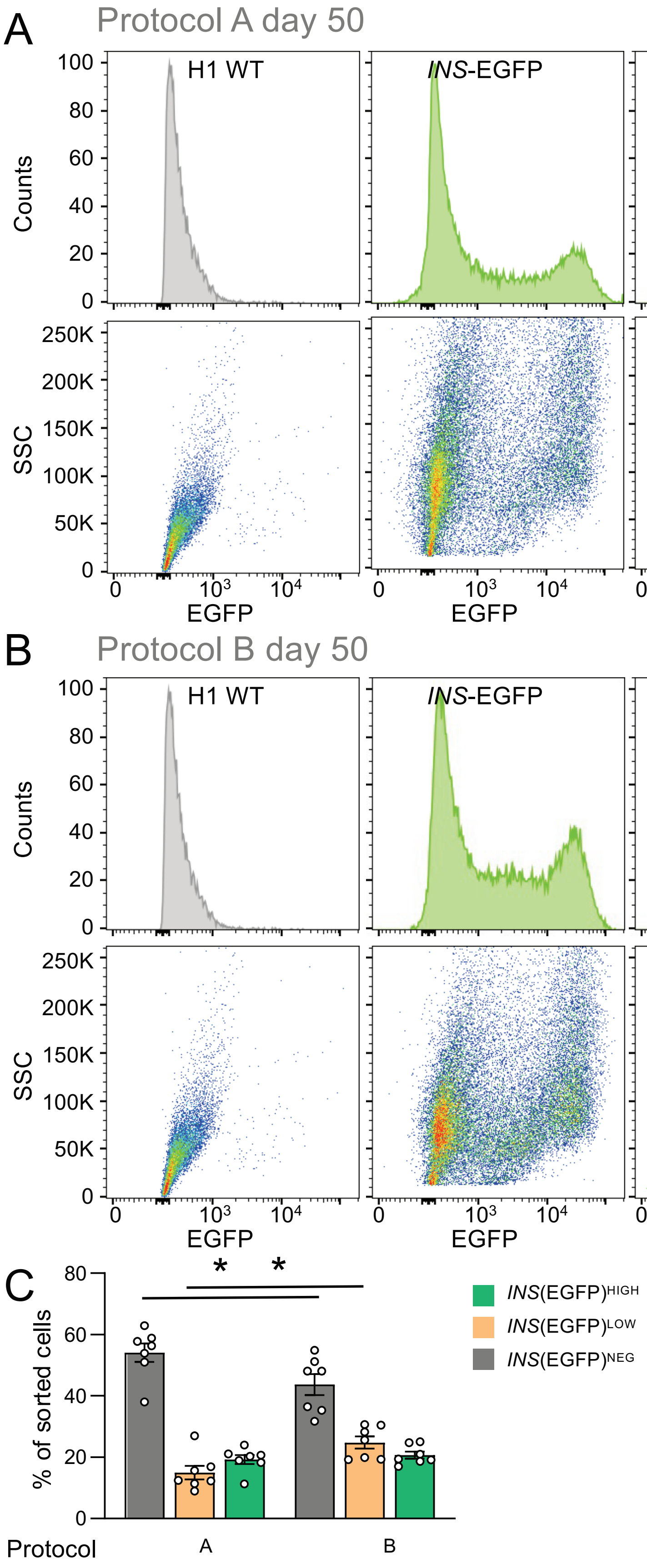


**Supplemental Figure 2. FACS analyses of *INS*-EGFP SCβ cells. (A,B)** FACS plots of stage 7 day 50 SCβ cells cultured with protocol A and B, respectively. **(C)** Cell state proportions of stage 7 day 50 SCβ cells cultured with protocol A and B. *$p < 0.05$

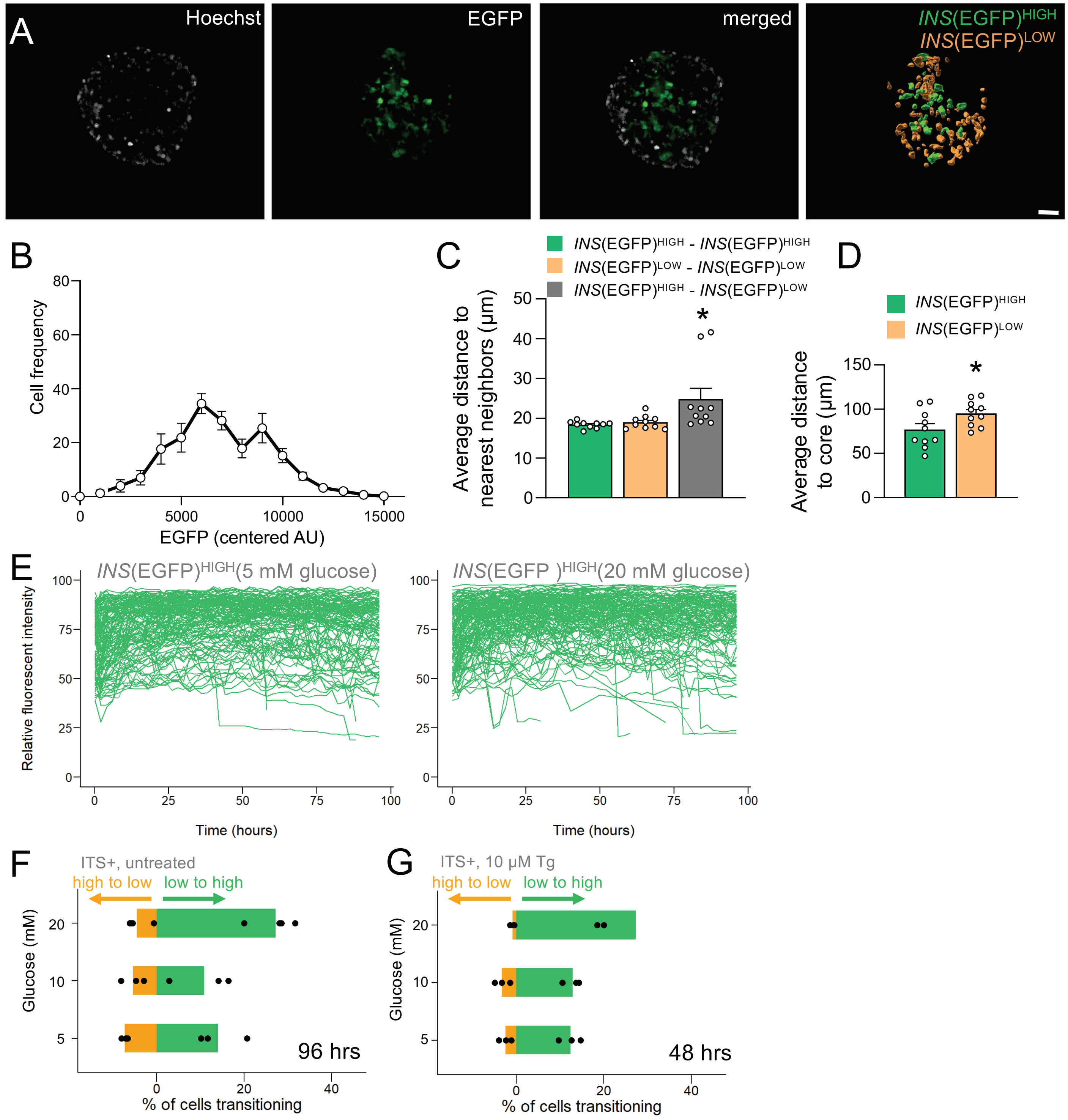


**Supplemental Figure 3. Dynamic insulin production in *INS*-EGFP SCβ cells. (A)** Example images of 3D live cell imaging of intact SCβ cell spheroids (n = 10 spheroids). Scale bar is 30 um. **(B)** Density plot of cells from intact *INS*-EGFP spheroids (n=10 spheroids). **(C)** Nearest neighbor analysis of cells cells from intact *INS*-EGFP spheroids. (n=10 spheroids). ANOVA. **(D)** Distance to core of cells cells from intact *INS*-EGFP spheroids. (n=10 spheroids). Student's T-test. **(E)** Example traces of cells which started experiments in the *INS*(EGFP)$^{HIGH}$ state. **(F)** *INS*-EGFP cell state transitions in media containing ITS but not treated with thapsigargin. **(G)** *INS*-EGFP cell state transitions in media containing ITS and treated with 10 μM thapsigargin. *p < 0.05.

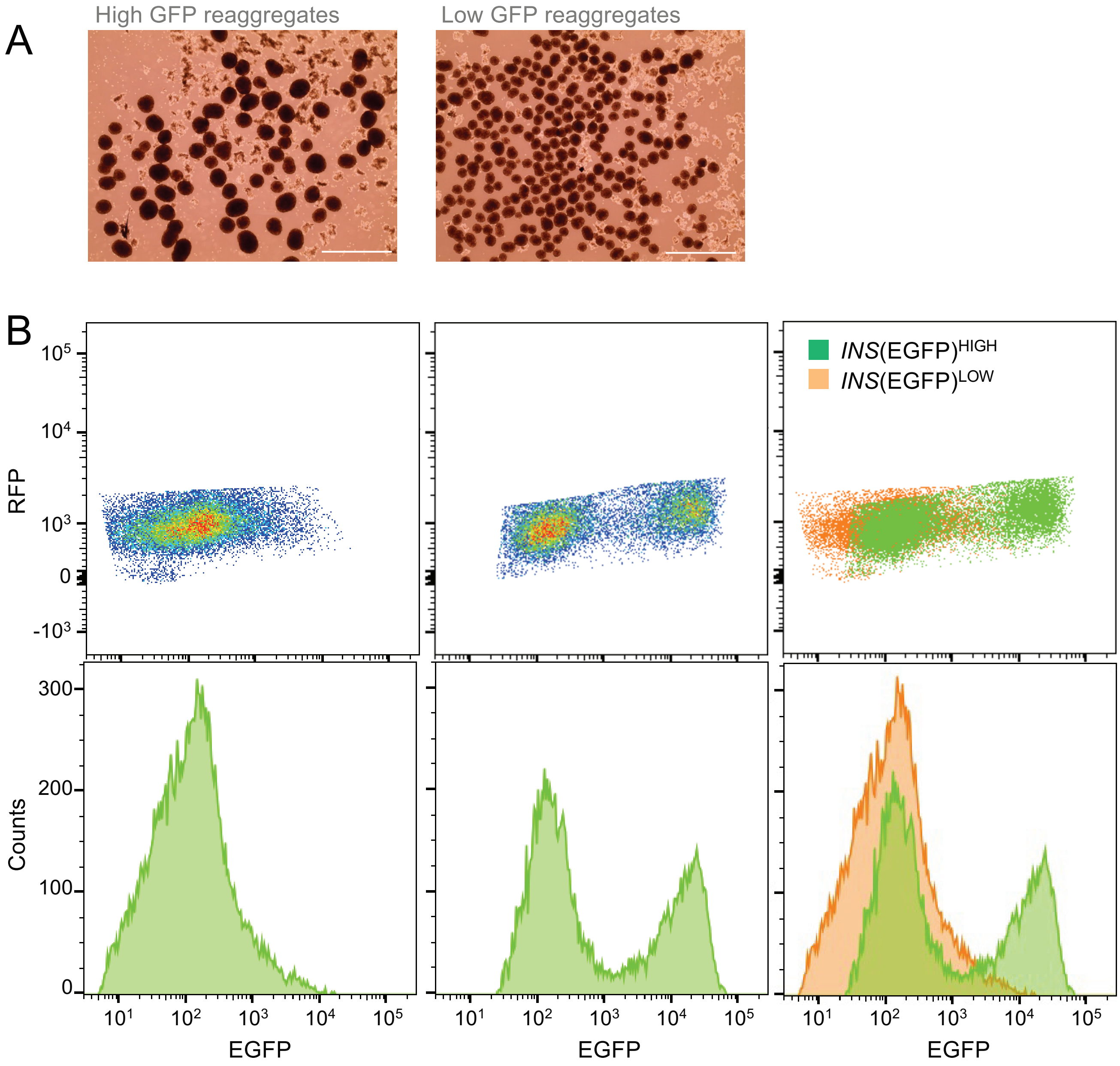


**Supplemental Figure 4. Reaggregation of FACS purified SCβ cells. (A)** Representative images of reaggregates made from FACS purified *INS*(EGFP)$^{HIGH}$ and *INS*(EGFP)$^{LOW}$ cells. Note the considerably larger size of the *INS*(EGFP)$^{HIGH}$ reaggregates. Scale bar is 1 mm. **(B)** Flow plots of EGFP fluorescence in *INS*(EGFP)$^{HIGH}$ and *INS*(EGFP)$^{LOW}$ reaggregates.

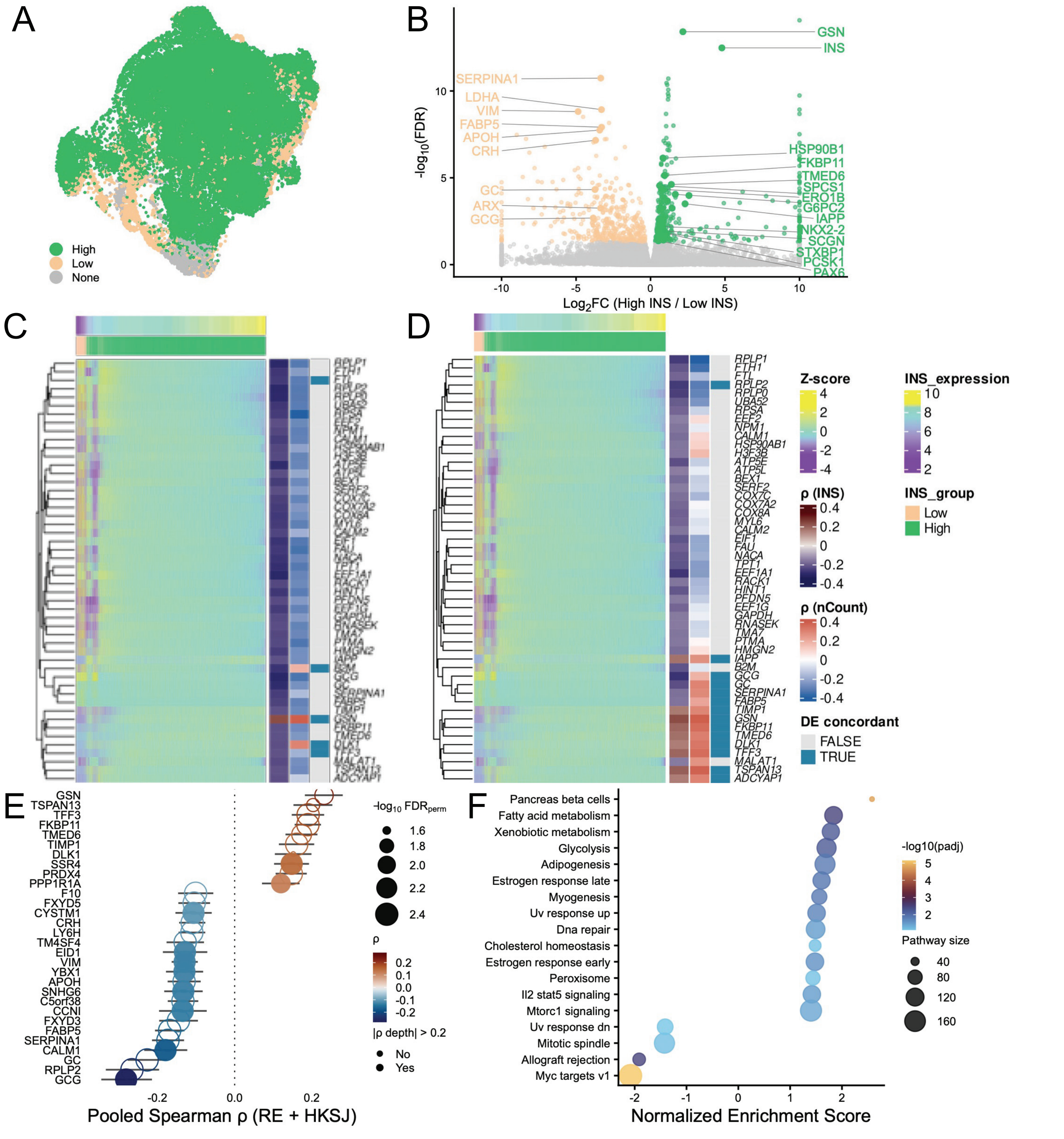


**Supplemental Figure 5. Differential gene expression and single-cell correlation analyses in high and low INS β cells.** **(A)** UMAP of non-diabetic β cells (n = 60,663 cells) colored by INS expression group assigned by per-donor Gaussian mixture model classification. **(B)** Volcano plot of pseudobulk differential expression (high vs low INS, depth-corrected model including donor and mean library size as covariates). **(C)** Heatmap of top 50 genes by |ρ| from within-donor Spearman correlation analysis, including ribosomal genes. Cells are ordered by INS expression. Row annotations show pooled Spearman ρ (INS), correlation with total transcript count ρ (nCount) as an indicator of potential depth confounding, and concordance with pseudobulk DE results. **(D)** As in (C), excluding ribosomal protein genes. **(E)** Lollipop plot of genes significant in both pseudobulk DE and correlation analyses (concordant hits). Open circles indicate genes with |ρ with nCount_RNA| > 0.2, suggesting potential library depth confounding. **(F)** GSEA of Hallmark pathways using within-donor Spearman correlation z-scores as the ranking metric. Shown are pathways significant (padj < 0.1) in either the DE-based or correlation-based GSEA.

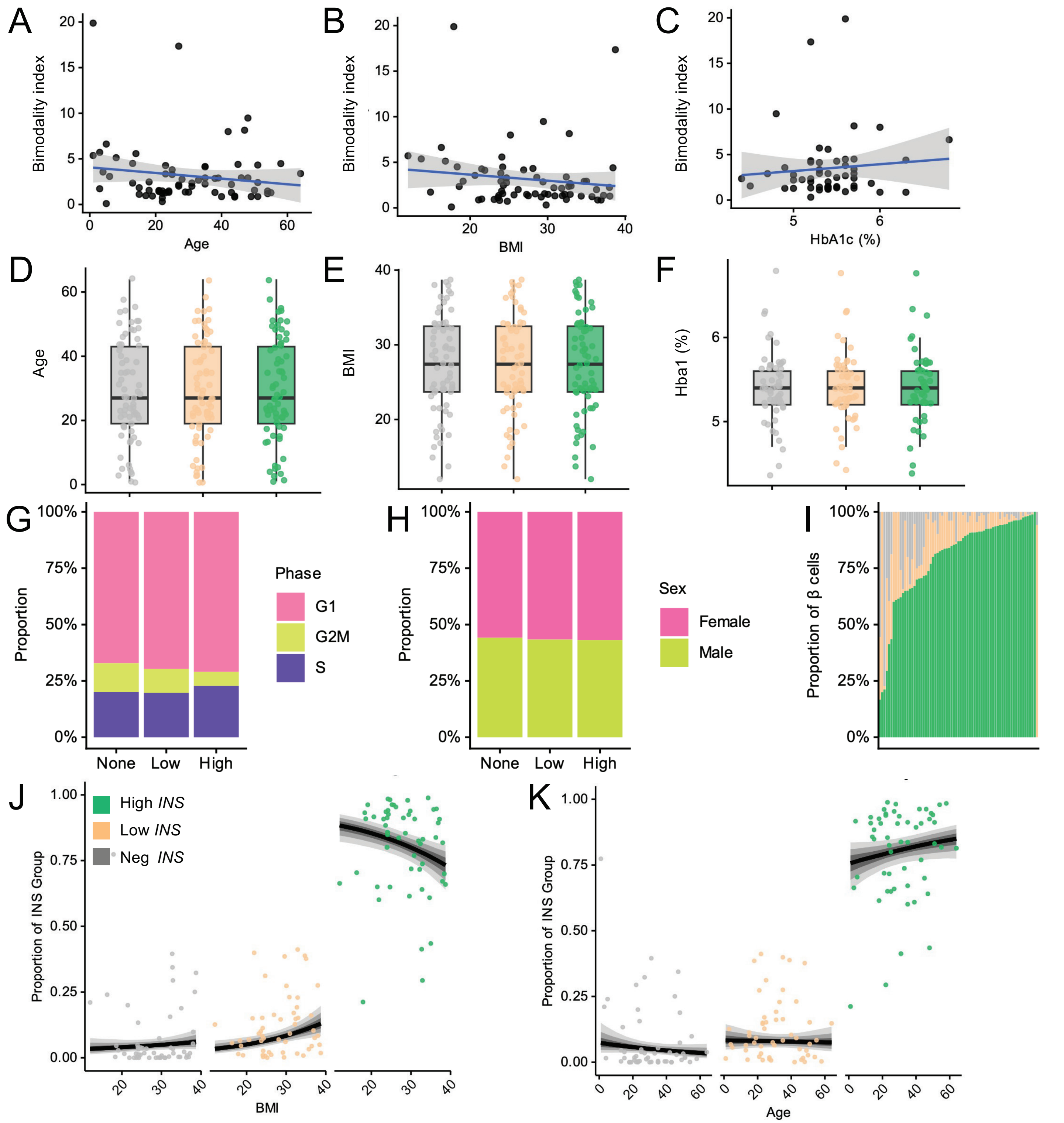


**Supplemental Figure 6. Characterization of INS mRNA expression groups in non-diabetic human β cells. (A–C)** Bimodality index of donor *INS* expression distributions plotted against donor age (A), BMI (B), and HbA1c (C). Each point represents one donor; the blue line shows a linear fit with 95% confidence interval. A modest negative correlation between BMI and bimodality index was observed (Spearman rho = −0.25, p = 0.065, donors with ≥50 cells). **(D–F)** Distribution of donor age (D), BMI (E), and HbA1c (F) across INS expression groups. Each point represents one donor; boxes show median and interquartile range. **(G)** Cell cycle phase proportions across *INS* groups. **(H)** Sex proportions across *INS* groups. **(I)** Proportion of β cells assigned to each *INS* group per donor (each bar represents one donor, n = 69), ordered by increasing proportion of High *INS* cells. **(J–K)** Posterior predicted proportions of cells in each *INS* group as a function of BMI (D) and age (E), derived from Bayesian zero-one-inflated beta regression models with study as a random effect. Shaded bands represent 50%, 80%, and 95% credible intervals. Points show observed donor-level proportions colored by *INS* group. LOO-IC model comparison did not identify BMI or age as convincingly superior to an intercept-only baseline model (all |ΔELPD| < 13, all SE > 4.5), indicating that neither variable reliably predicted *INS* group proportions across donors.

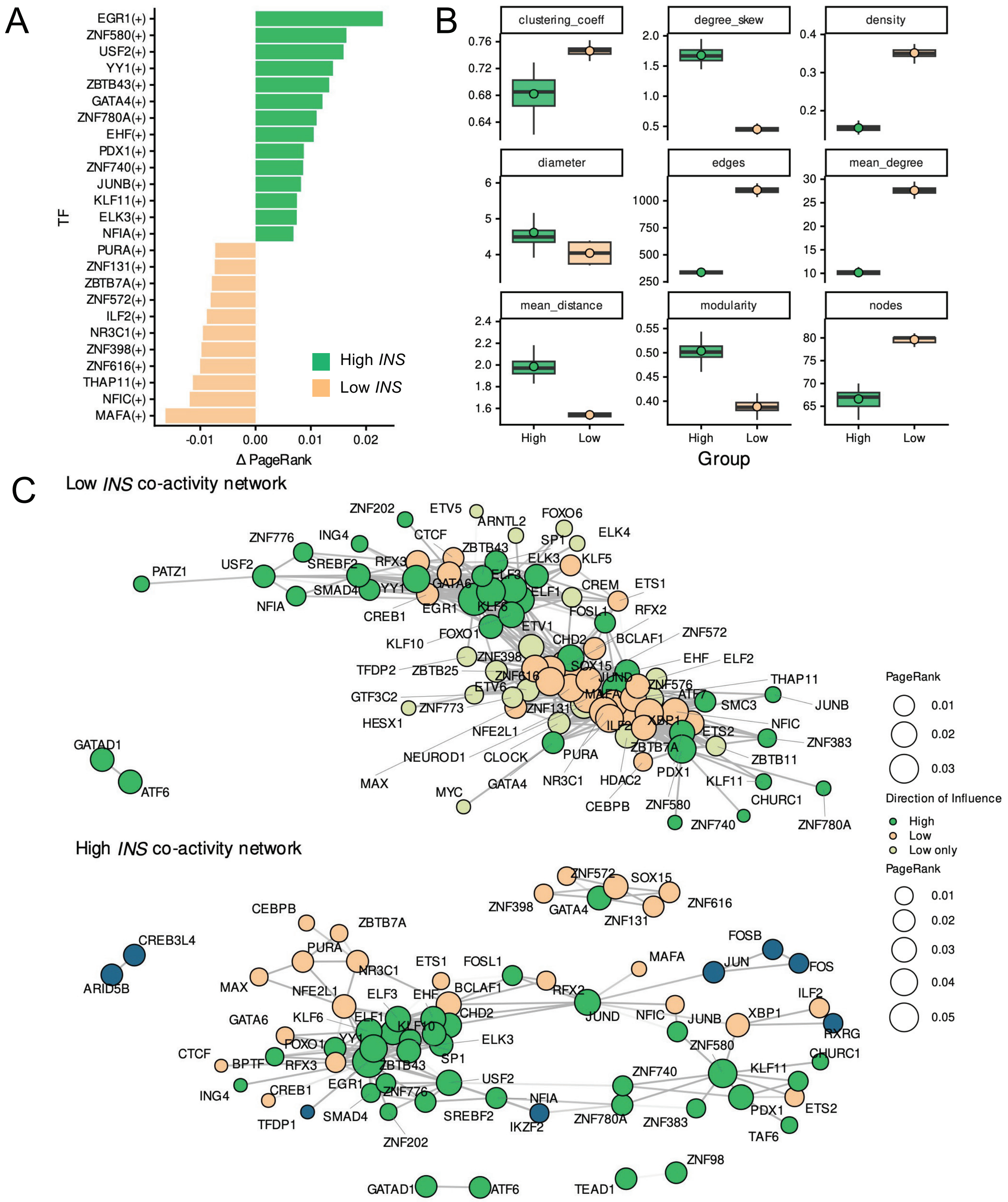


**Supplemental Figure 7. Quantitative characterization of transcription factor co-activity networks in high and low INS β cells. (A)** Change in mean PageRank centrality (ΔPageRank = High − Low) for transcription factors present in both networks. Positive values indicate greater centrality in high *INS* networks. **(B)** Bootstrap distributions of network topology metrics for high and low *INS* co-activity networks across 100 subsampled iterations. High *INS* networks show higher clustering coefficient and modularity with lower mean degree, consistent with a more community-structured topology. Points indicate bootstrap means. **(C–D)** Fully labeled co-activity networks for low *INS* (C) and high *INS* (D) β cells. Node size reflects mean PageRank centrality; node color indicates direction of influence. Edge opacity reflects the proportion of bootstrap iterations in which the co-activity edge was detected (threshold: ≥50 of 100 iterations).

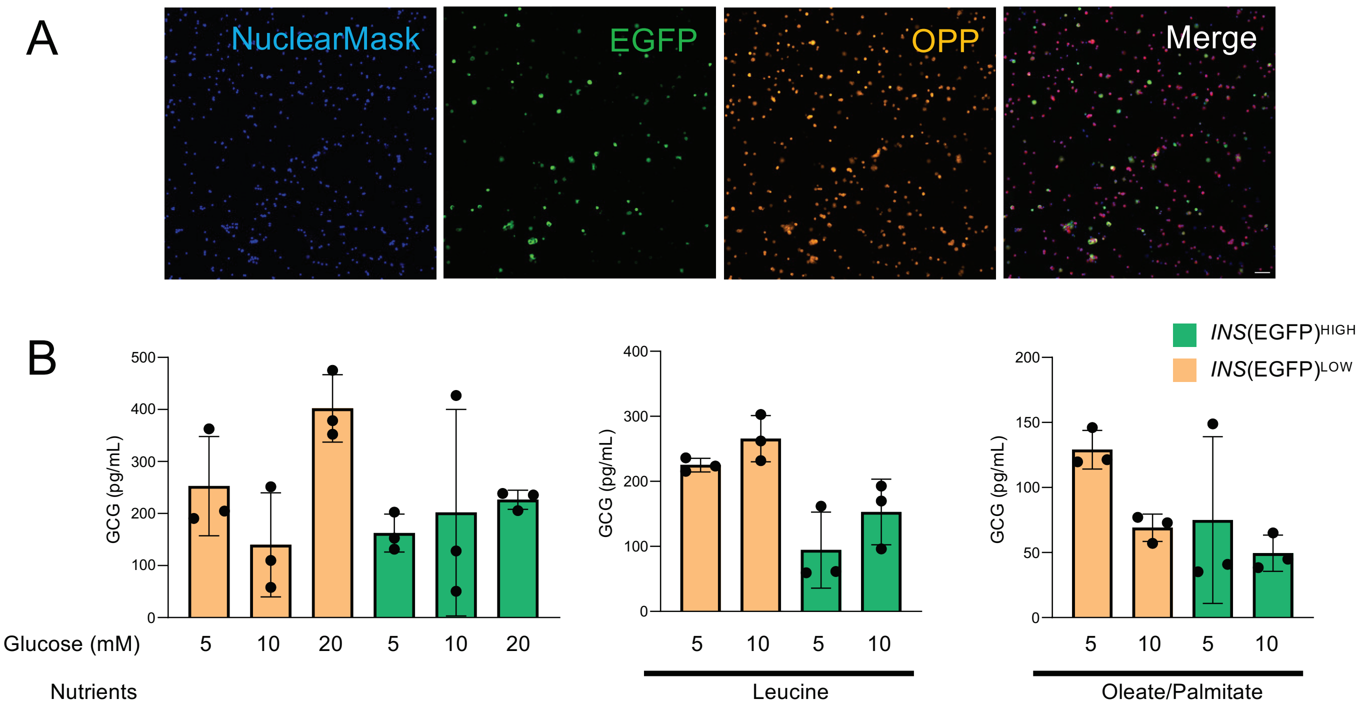


**Supplemental Figure 8. Functional analyses of *INS*(EGFP)$^{HIGH}$ and *INS*(EGFP)$^{LOW}$ cells. (A)** Representative images of the OPP assay. **(B)** Glucose stimulated glucagon secretion of SCβ cells cells sorted into the high and low *INS* states. (n=3 differentiations).

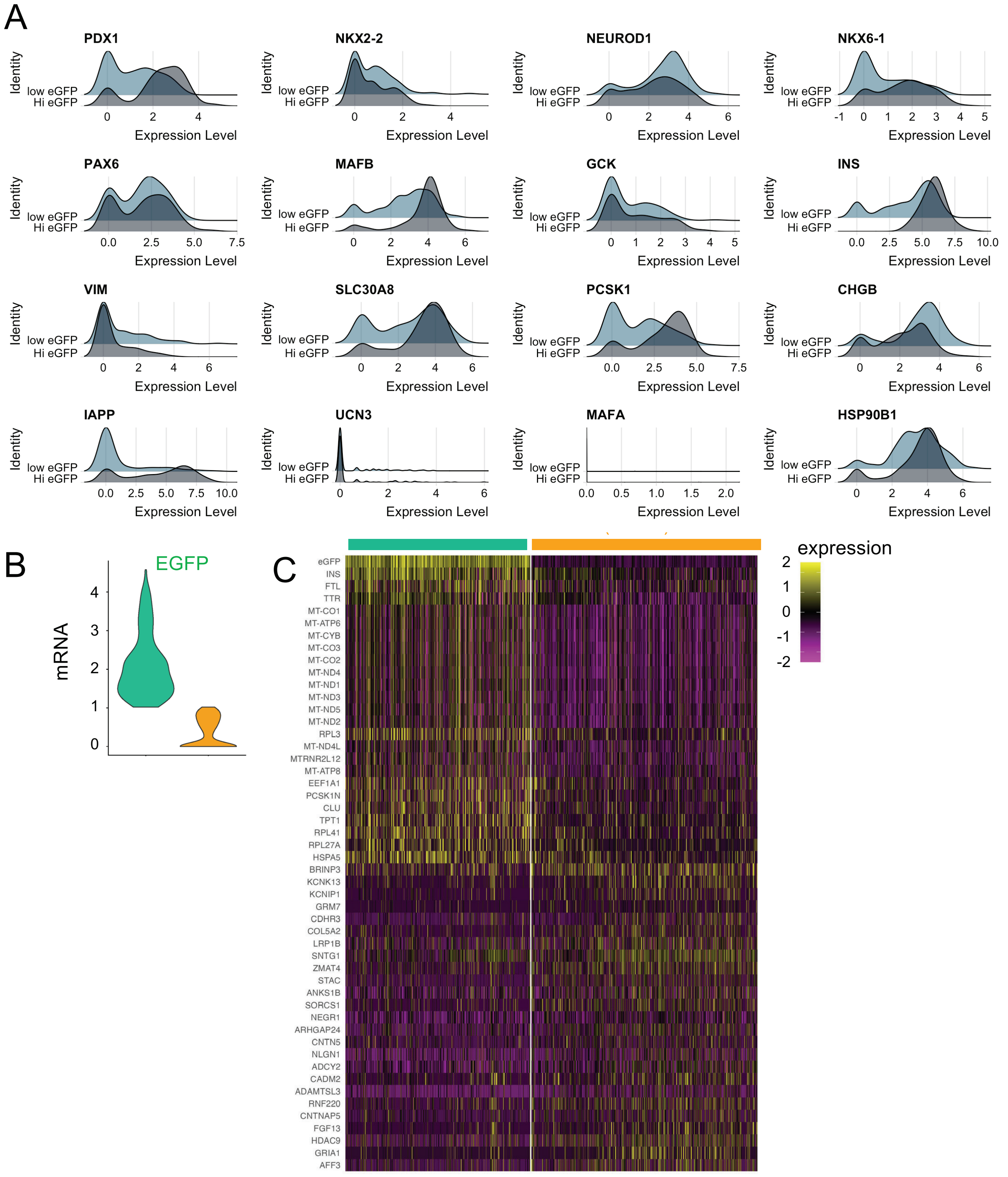


**Supplemental Figure 9. Categorical scRNAseq comparison of *INS*(EGFP)$^{HIGH}$ and *INS*(EGFP)$^{LOW}$ SCβ cells based on EGFP levels. (A)** Patch-seq data of β cell and ER stress markers in *INS*(EGFP)$^{HIGH}$ and *INS*(EGFP)$^{LOW}$ cells. **(B)** *Egfp* mRNA distribution in *INS*(EGFP)$^{HIGH}$ versus *INS*(EGFP)$^{LOW}$ cells. **(C)** Heatmap of categorical differential gene analysis of *INS*(EGFP)$^{HIGH}$ and *INS*(EGFP)$^{LOW}$ cells.

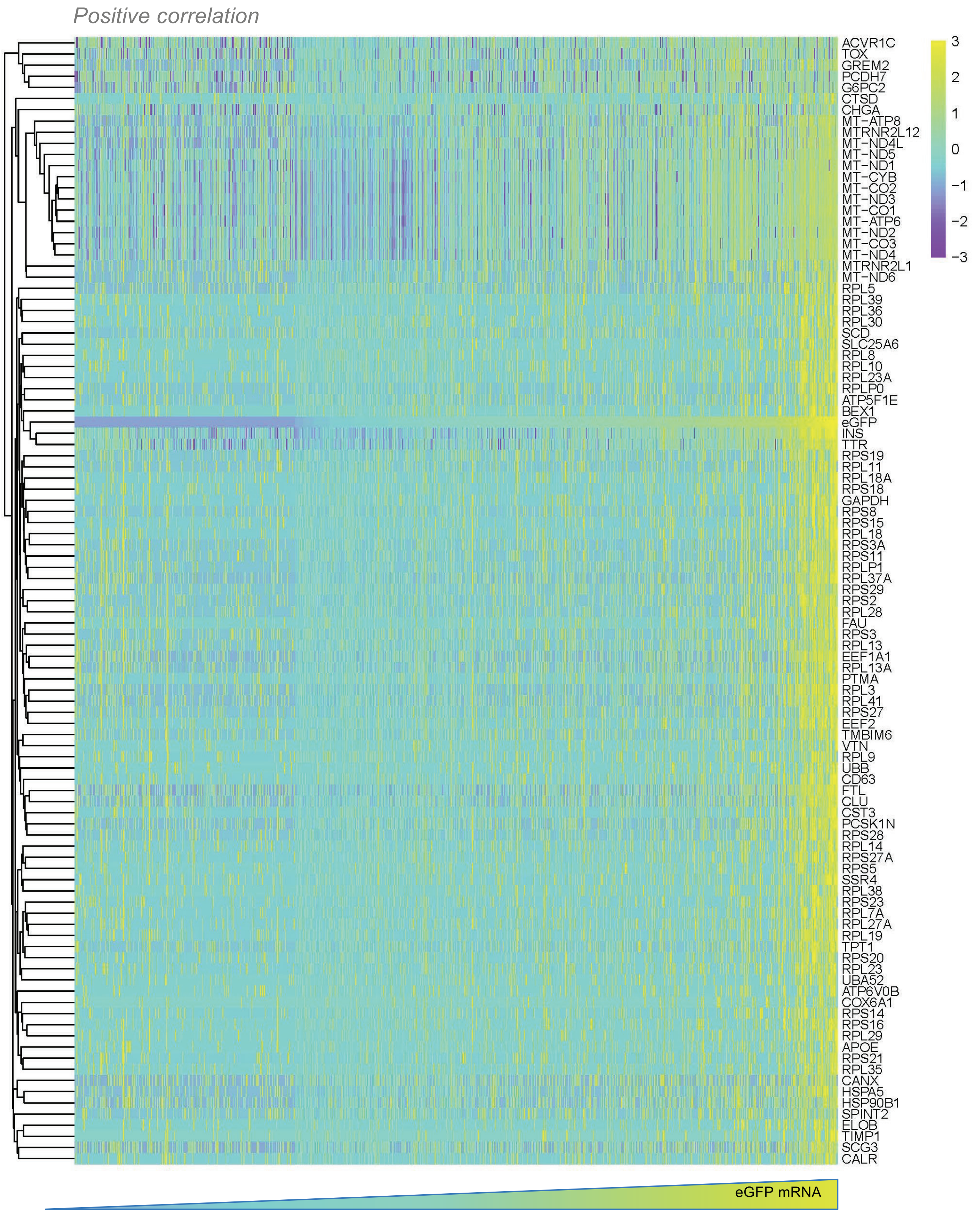


**Supplemental Figure 10. Top 100 mRNAs that positively correlate with EGFP in SCβ cells.**

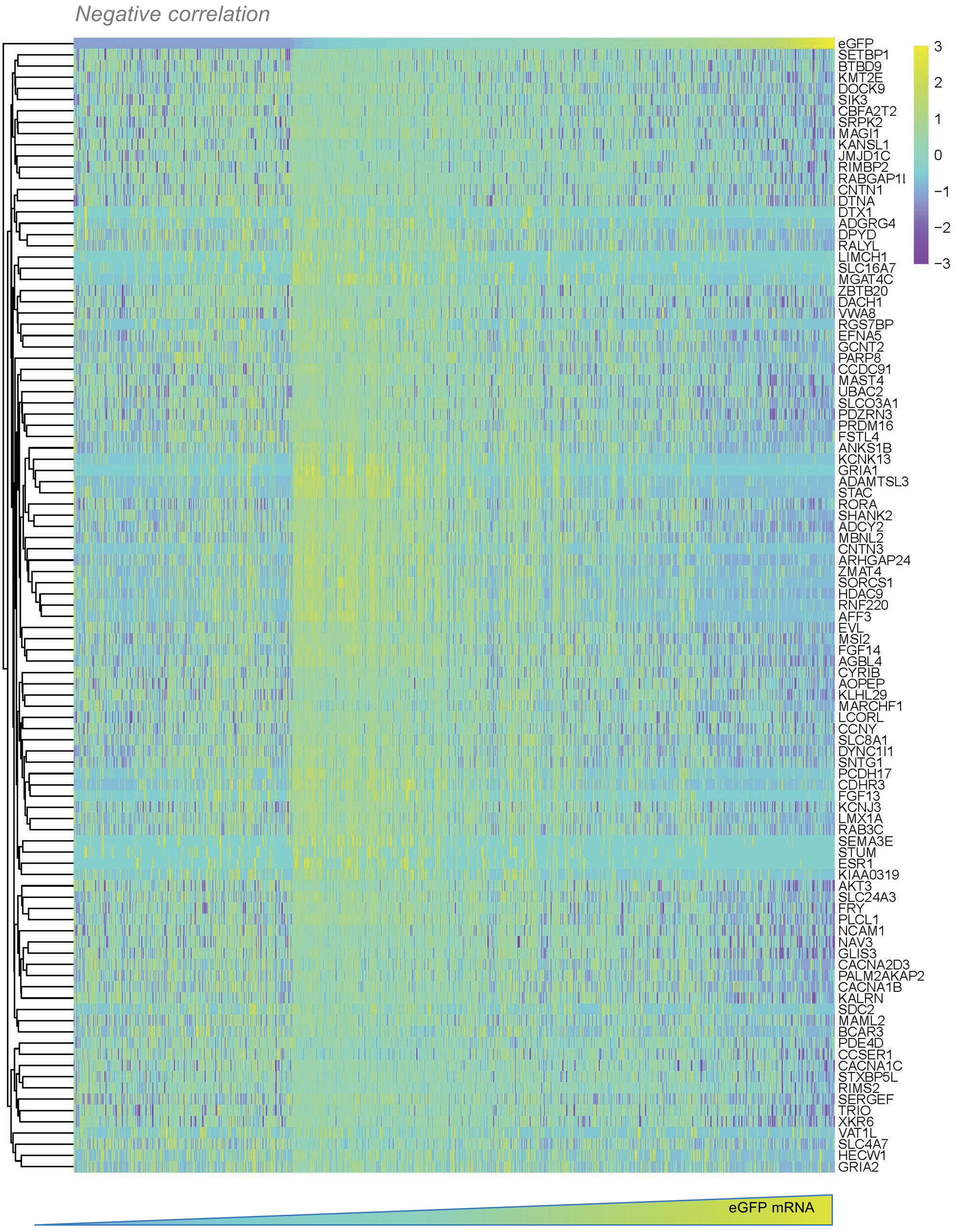


**Supplemental Figure 11. Top 100 mRNAs that negatively correlate with EGFP in SCβ cells.**

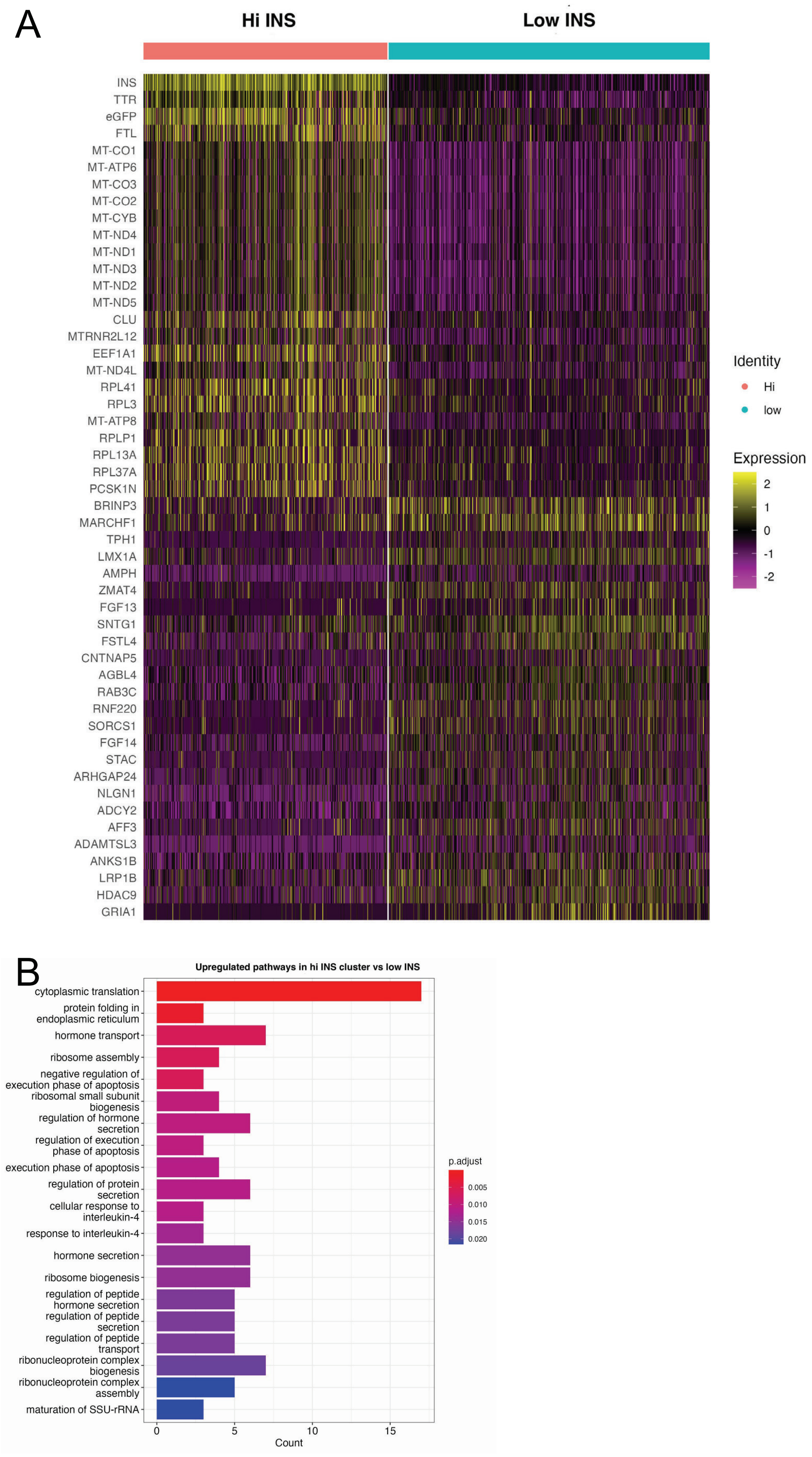


**Supplemental Figure 12. Categorical scRNAseq comparison of *INS*(EGFP)$^{HIGH}$ and *INS*(EGFP)$^{LOW}$ SCβ cells based on *INS* levels. (A)** Differential gene analysis of cells with high and low *INS* mRNA. **(B)** Overrepresentation analysis reveals pathways enriched in cells with high *INS* mRNA.

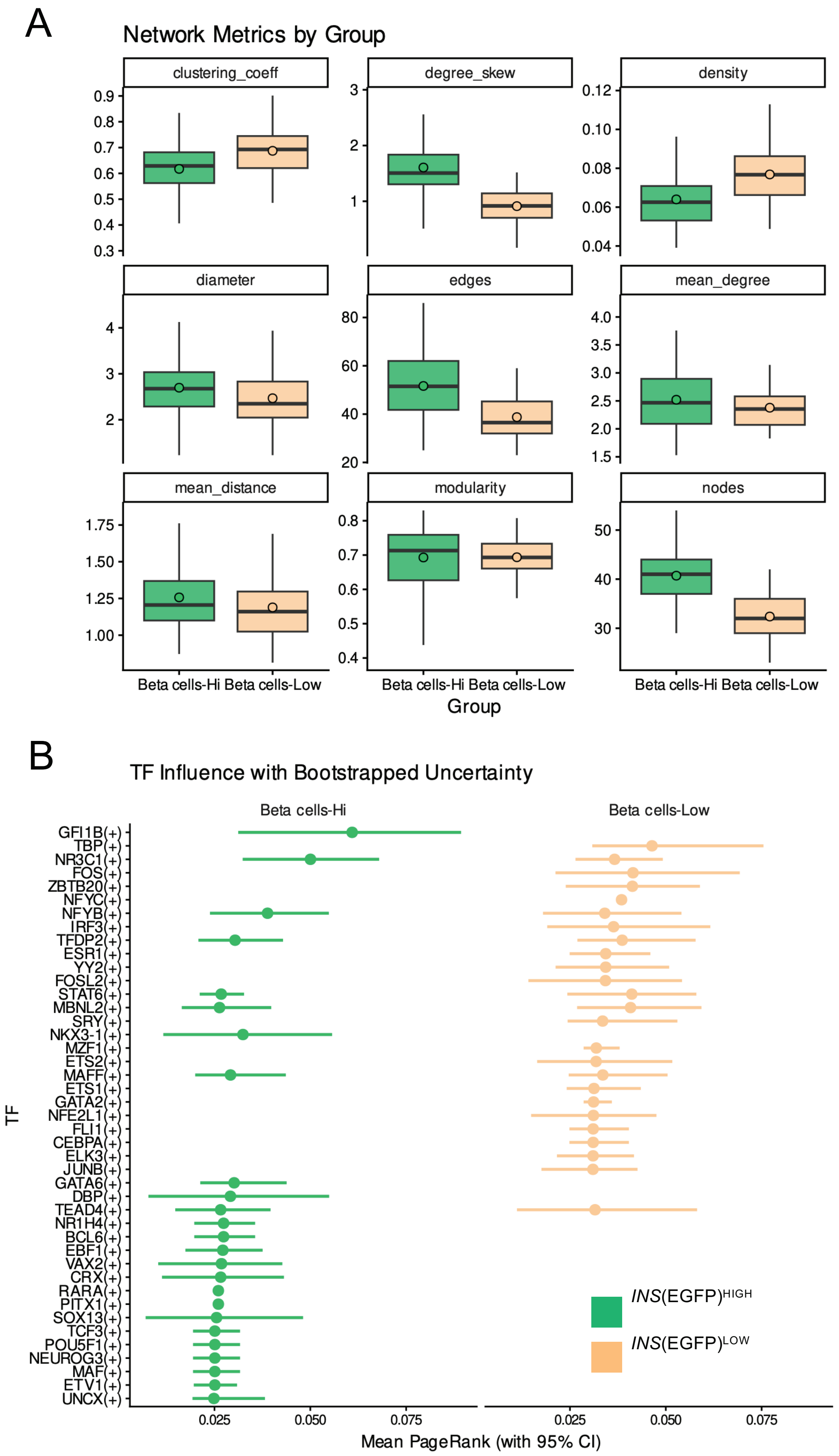


**Supplemental Figure 13. SCENIC and patch-seq analyses of *INS*(EGFP)$^{HIGH}$ and *INS*(EGFP)$^{LOW}$ SCβ cells. (A)** Quantification of network analysis metrics. **(B)** Top TFs ranked by influence scores in *INS*(EGFP)$^{HIGH}$ and *INS*(EGFP)$^{LOW}$ states.

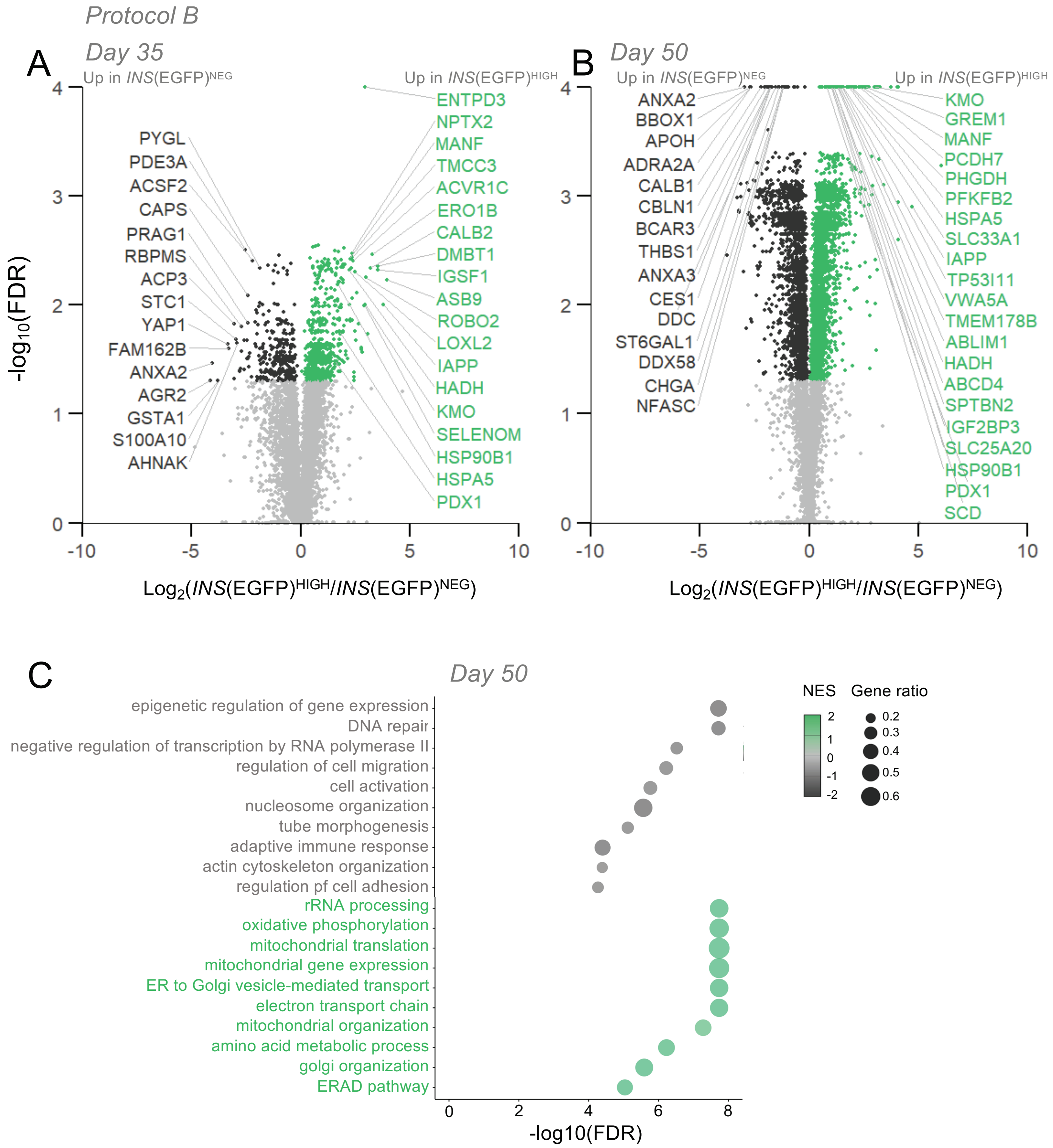


**Supplemental Figure 14. Proteomic analyses of *INS*(EGFP)$^{HIGH}$ and *INS*(EGFP)$^{NEG}$ SCβ cells cultured using protocol B. (A-B)** Volcano plots depicting differentially abundant proteins in *INS*(EGFP)$^{HIGH}$ and *INS*(EGFP)$^{NEG}$ SCβ cells at S7 day 35 and 50**. (C)** Gene set enrichment analysis revealing upregulated pathways in *INS*(EGFP)$^{HIGH}$ and *INS*(EGFP)$^{NEG}$ SCβ cells at S7 day 50**.**

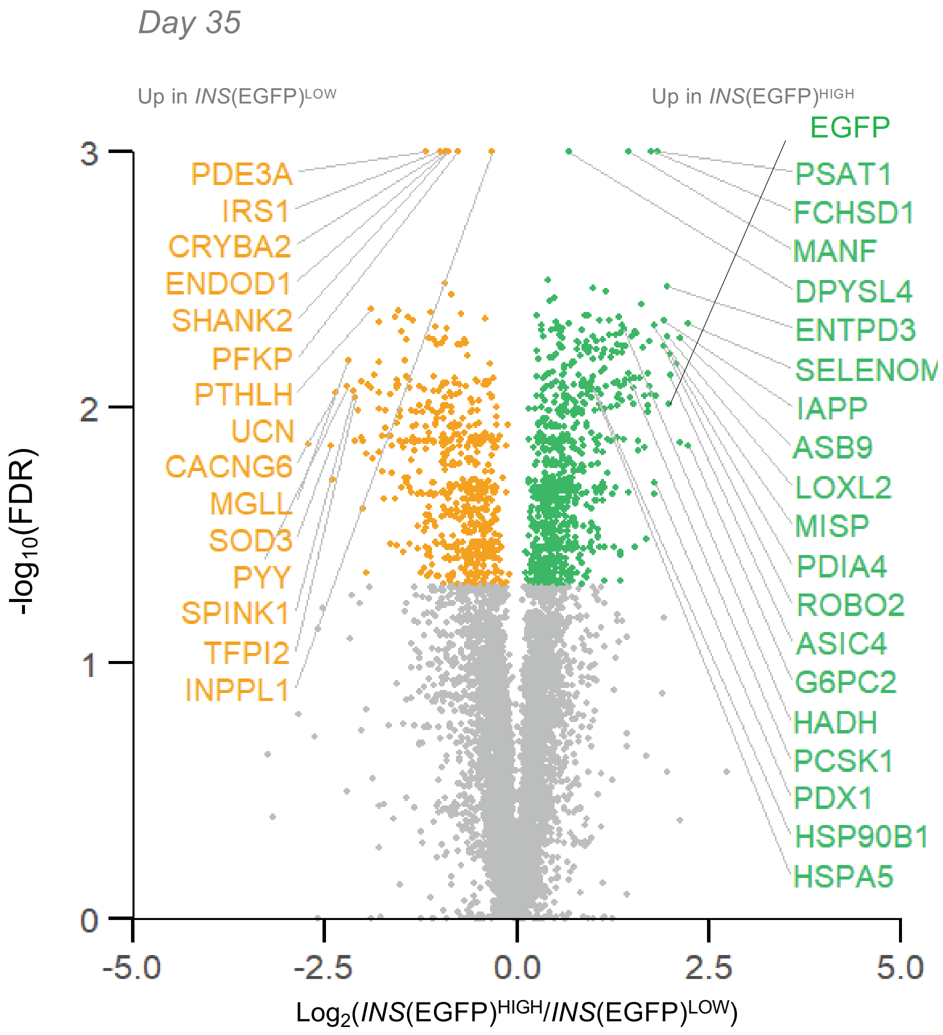


**Supplemental Figure 15.** Volcano plot depicting differentially abundant proteins in *INS*(EGFP)$^{HIGH}$ and *INS*(EGFP)$^{LOW}$ SCβ cells at S7 day 35 using protocol B.

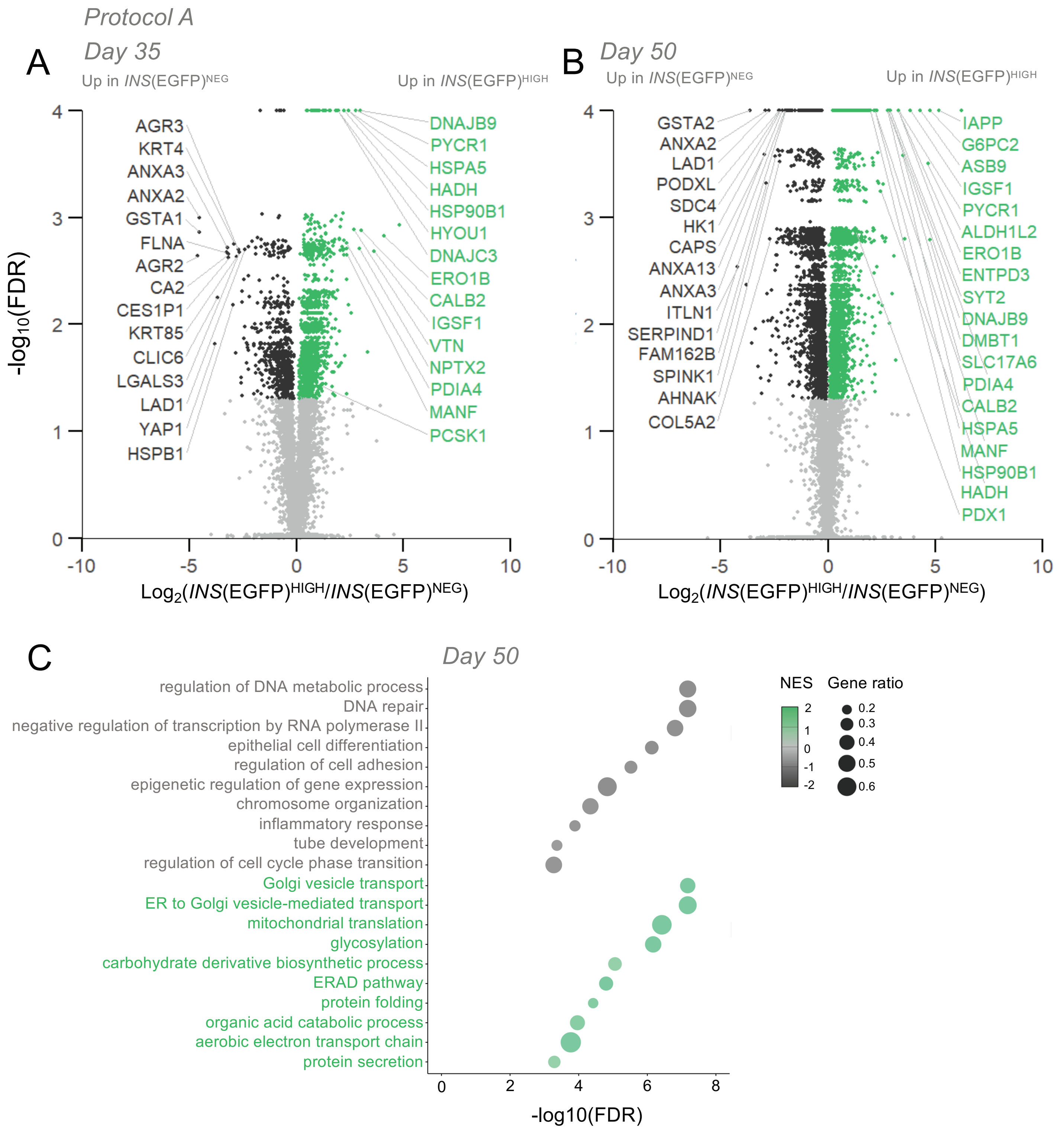


**Supplemental Figure 16. Proteomic analyses of *INS*(EGFP)$^{HIGH}$ and *INS*(EGFP)$^{NEG}$ SCβ cells cultured using protocol A. (A-B)** Volcano plots depicting differentially abundant proteins in *INS*(EGFP)$^{HIGH}$ and *INS*(EGFP)$^{NEG}$ SCβ cells at S7 day 35 and 50**. (C)** Gene set enrichment analysis revealing upregulated pathways in *INS*(EGFP)$^{HIGH}$ and *INS*(EGFP)$^{NEG}$ SCβ cells at S7 day 50**.**

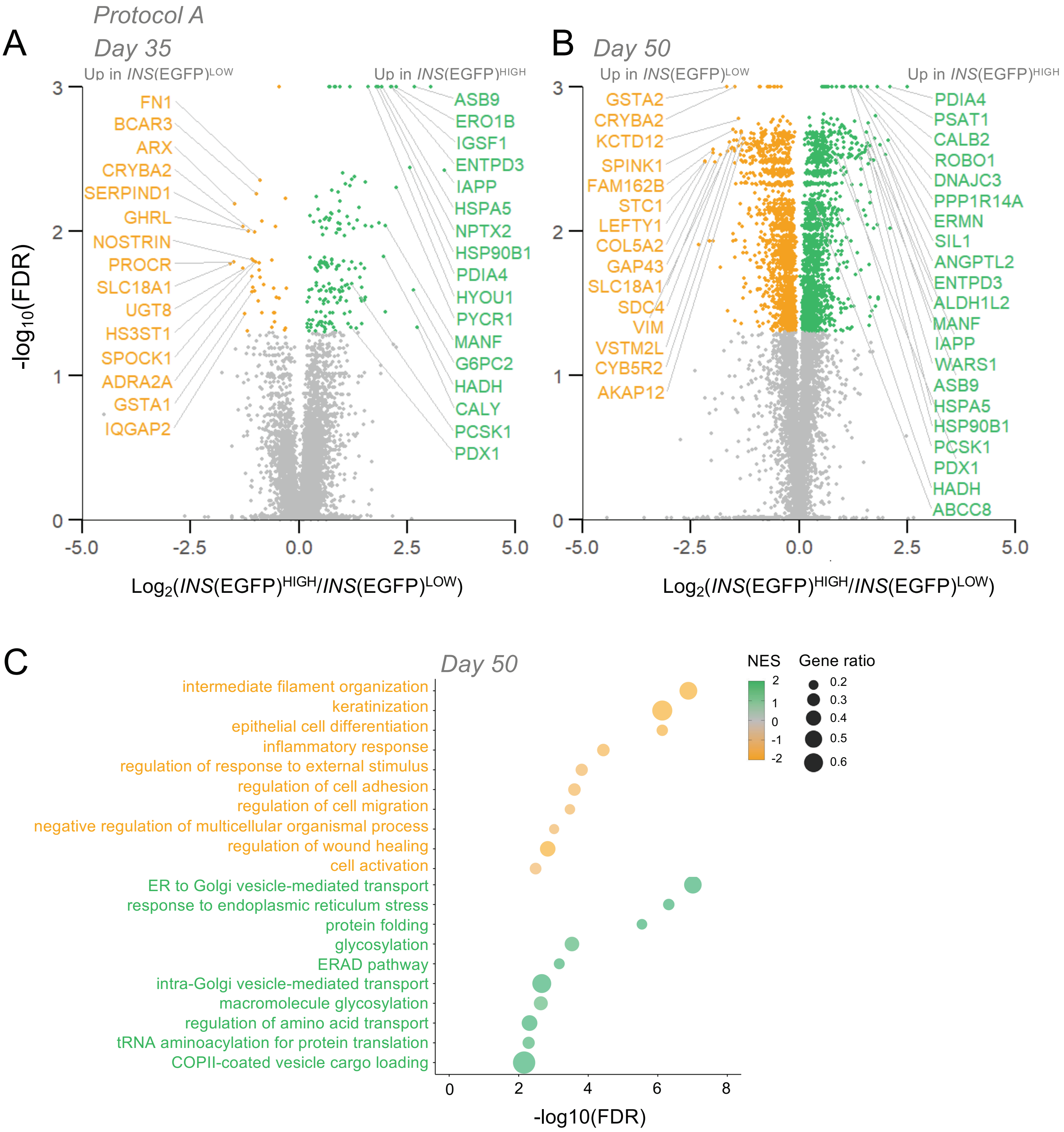


**Supplemental Figure 17. Proteomic analyses of *INS*(EGFP)$^{HIGH}$ and *INS*(EGFP)$^{LOW}$ SCβ cells cultured using protocol A. (A-B)** Volcano plots depicting differentially abundant proteins in *INS*(EGFP)$^{HIGH}$ and *INS*(EGFP)$^{LOW}$ SCβ cells at S7 day 35 and 50**. (C)** Gene set enrichment analysis revealing upregulated pathways in *INS*(EGFP)$^{HIGH}$ and *INS*(EGFP)$^{LOW}$ SCβ cells at S7 day 50**.**

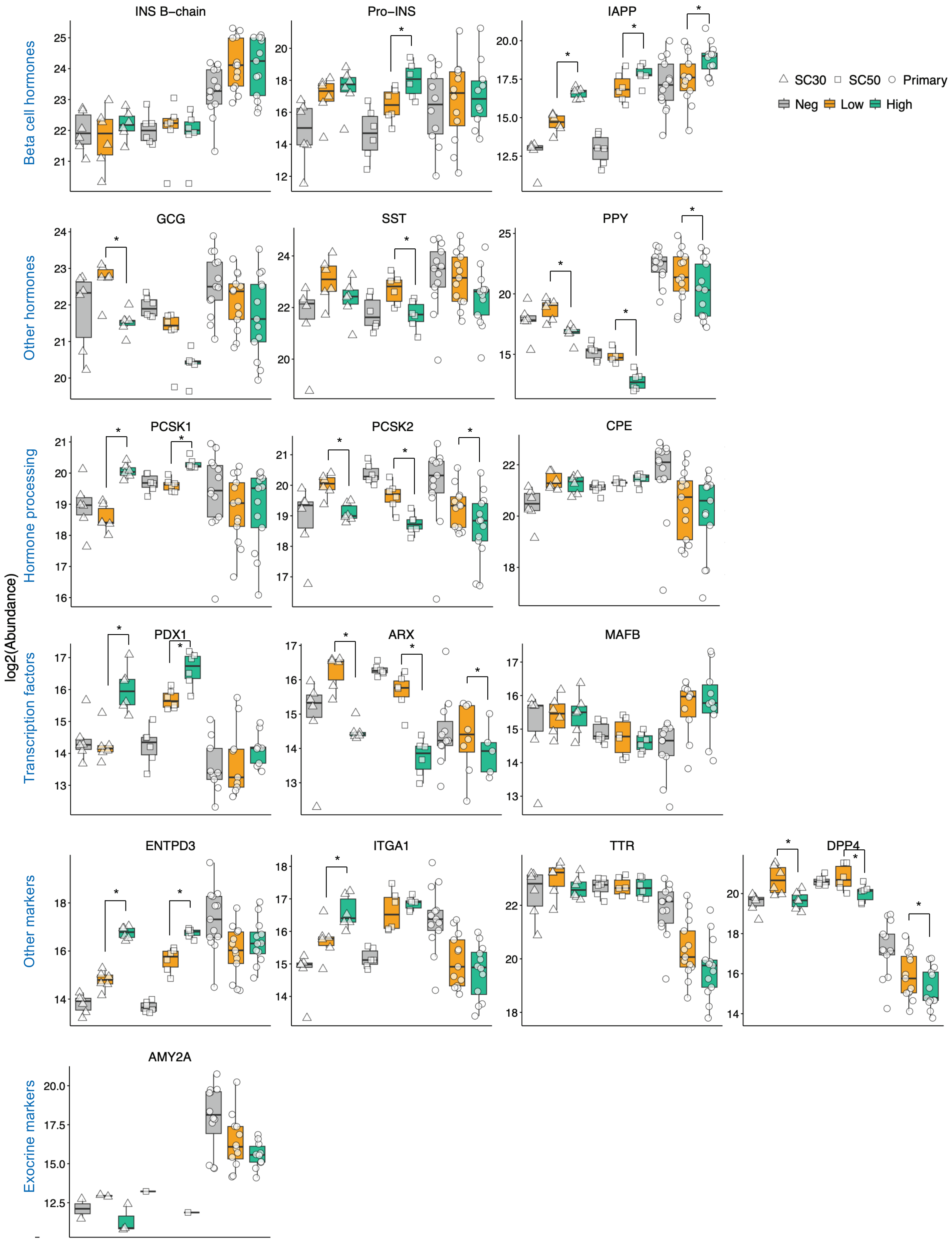


**Supplemental Figure 18. Protein abundance of key islet and pancreas cell markers across proteomics datasets.** Box plots of key proteins. * Multiple paired t-tests, q < 0.05.

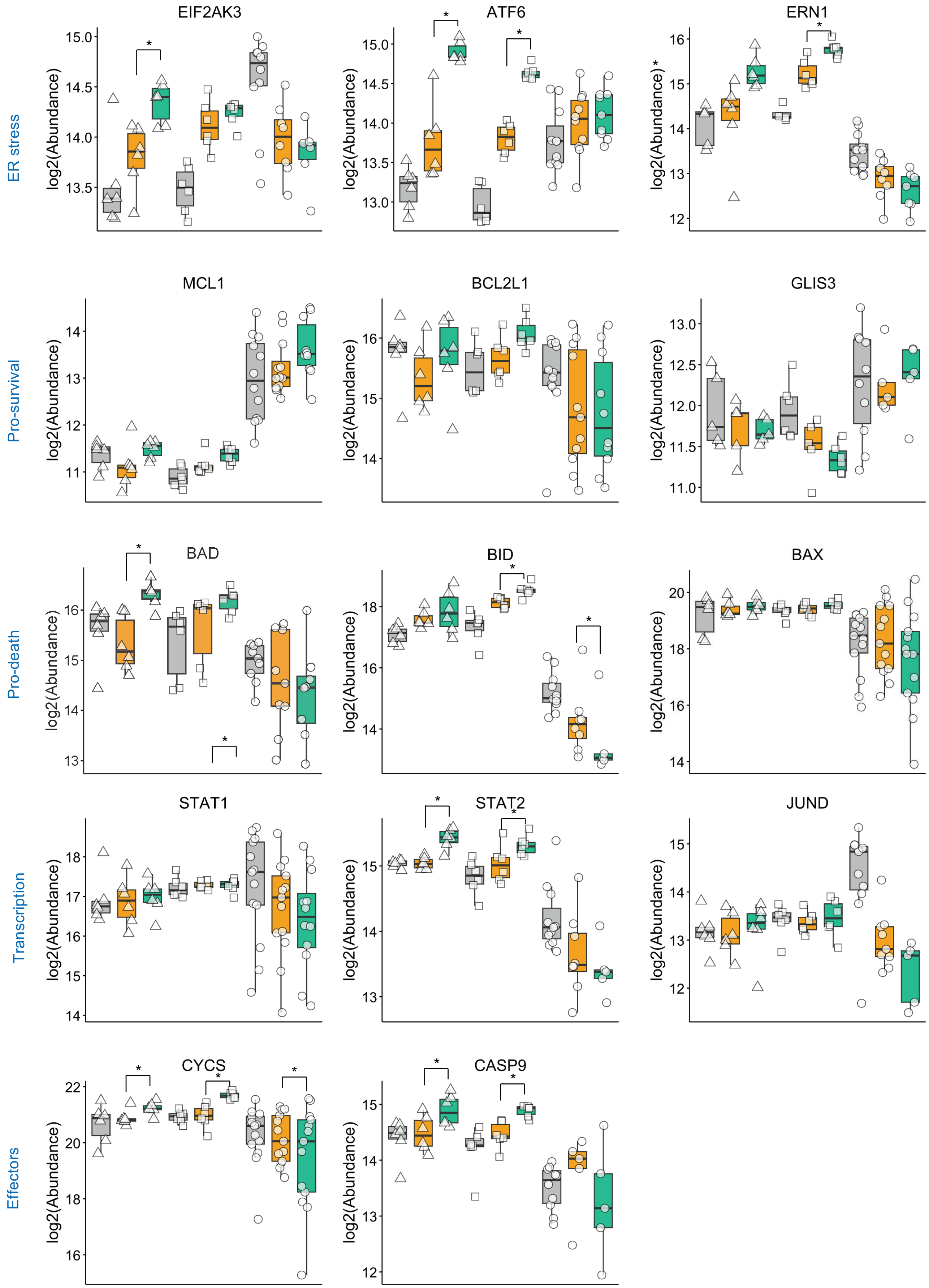


**Supplemental Figure 19. Protein abundance of key cell death markers in INS low and INS high beta cells across proteomics datasets.** Boxplots of key proteins. * Multiple paired t-tests, q < 0.05.

### Supplemental Tables

#### Table S1. Human islet donor metadata.

| Donor ID | RRID | HLA A2 | Diagnosis | adjusted diabetes status | Age (years) | Sex | HbA1c (%) | BMI ($kg/m^2$) | Donation Type | Experiments |
|---|---|---|---|---|---|---|---|---|---|---|
| **R541** | | positive | n/a | Pre-T2D | 66 | F | 6.10% | 26.4 | NDD Neurological | proteomics |
| **R543** | | positive | n/a | n/a | 54 | F | 5.40% | 20.2 | NDD Neurological | proteomics |
| **R559** | | positive | n/a | n/a | 64 | F | 5.60% | 31.7 | NDD Neurological | proteomics |
| **R562** | | negative | n/a | T2D | 45 | F | 7.10% | 30.9 | NDD Neurological | proteomics |
| **R563** | | positive | n/a | n/a | 59 | M | 4.50% | 26.6 | NDD Neurological | proteomics |
| **R564** | | negative | n/a | n/a | 37 | M | 5.40% | 28.7 | NDD Neurological | proteomics |
| **R568** | | positive | n/a | n/a | 60 | M | 5.50% | 21.1 | NDD Neurological | proteomics |
| **R570** | | negative | n/a | n/a | 45 | F | 5.40% | 31.3 | NDD Neurological | proteomics |
| **R574** | | positive | n/a | n/a | 45 | M | 5.10% | 25 | NDD Neurological | proteomics |
| **R576** | | positive | n/a | Pre-T2D | 38 | M | 5.80% | 32.5 | NDD Neurological | proteomics |
| **R583** | | positive | n/a | n/a | 33 | M | 5.70% | 23.1 | DNC | proteomics, cell death assay |
| **R585** | | positive | T2D | T2D | 66 | F | 5.80% | 31.5 | DNC | proteomics |
| **R589** | | negative | n/a | n/a | 55 | M | 5.40% | 24.2 | DCC | Cell death assay |
| **R592** | | positive | n/a | n/a | 48 | M | 5.50% | 33.2 | DCC | proteomics, cell death assay |
| **R595** | | negative | n/a | n/a | 57 | M | 5.50% | 33.1 | MAiD | OPP assay |

**Table S2. Key resources**

| Reagent/resource | Designation | Source/reference | RRID |
|---|---|---|---|
| Francis Lynn lab | *INS*-EGFP cell line | | N/A |
| Reagent | RPMI 1640 | Invitrogen, catalog number 11875 | N/A |
| Reagent | Fetal bovine serum | Invitrogen, catalog number 12483020 | N/A |
| Software | GraphPad Prism | GraphPad Software | N/A |
| Software | ImageXpress | Molecular Devices | N/A |
| Software | R | R Foundation for Statistical Computing | N/A |

**Table S3. DIA windows**

| #MS Type | Cycle Id | Start IM [1/K0] | End IM [1/K0] | Start Mass [m/z] | End Mass [m/z] | CE [eV] |
|---|---|---|---|---|---|---|
| **MS1** | 0 | - | - | - | - | - |
| **PASEF** | 1 | 0.6 | 0.805 | 350.68 | 382.04 | - |
| **PASEF** | 1 | 0.805 | 0.965 | 559.8 | 567.5 | - |
| **PASEF** | 1 | 0.965 | 1.12 | 725.13 | 736.24 | - |
| **PASEF** | 2 | 0.6 | 0.87 | 412.39 | 425.4 | - |
| **PASEF** | 2 | 0.87 | 0.977 | 574.21 | 582.01 | - |
| **PASEF** | 2 | 0.977 | 1.15 | 746.34 | 758.38 | - |
| **PASEF** | 3 | 0.6 | 0.89 | 437.42 | 447.23 | - |
| **PASEF** | 3 | 0.89 | 0.986 | 588.81 | 596.81 | - |
| **PASEF** | 3 | 0.986 | 1.19 | 769.41 | 782.64 | - |
| **PASEF** | 4 | 0.6 | 0.9 | 456.05 | 464.9 | - |
| **PASEF** | 4 | 0.9 | 0.995 | 603.8 | 612.07 | - |
| **PASEF** | 4 | 0.995 | 1.23 | 794.87 | 808.65 | - |
| **PASEF** | 5 | 0.6 | 0.906 | 472.76 | 480.93 | - |
| **PASEF** | 5 | 0.906 | 1.006 | 619.33 | 627.84 | - |
| **PASEF** | 5 | 1.006 | 1.27 | 821.43 | 837.18 | - |
| **PASEF** | 6 | 0.6 | 0.91 | 488.1 | 495.93 | - |
| **PASEF** | 6 | 0.91 | 1.025 | 635.35 | 644.1 | - |
| **PASEF** | 6 | 1.025 | 1.31 | 851.93 | 869.71 | - |
| **PASEF** | 7 | 0.6 | 0.917 | 502.77 | 510.43 | - |
| **PASEF** | 7 | 0.917 | 1.045 | 651.86 | 660.85 | - |
| **PASEF** | 7 | 1.045 | 1.35 | 886.49 | 908.17 | - |
| **PASEF** | 8 | 0.6 | 0.927 | 517.08 | 524.68 | - |

| | | | | | | |
|---|---|---|---|---|---|---|
| **PASEF** | 8 | 0.927 | 1.064 | 668.83 | 678.09 | - |
| **PASEF** | 8 | 1.064 | 1.39 | 928.86 | 956.26 | - |
| **PASEF** | 9 | 0.6 | 0.94 | 531.29 | 539.03 | - |
| **PASEF** | 9 | 0.94 | 1.085 | 686.35 | 696.12 | - |
| **PASEF** | 9 | 1.085 | 1.43 | 982.67 | 1021.61 | - |
| **PASEF** | 10 | 0.6 | 0.955 | 545.77 | 553.28 | - |
| **PASEF** | 10 | 0.955 | 1.105 | 704.89 | 715.51 | - |
| **PASEF** | 10 | 1.105 | 1.45 | 1059.54 | 1155.34 | - |
| **PASEF** | 11 | 0.6 | 0.85 | 381.54 | 412.89 | - |
| **PASEF** | 11 | 0.85 | 0.97 | 567.01 | 574.71 | - |
| **PASEF** | 11 | 0.97 | 1.13 | 735.74 | 746.84 | - |
| **PASEF** | 12 | 0.6 | 0.884 | 424.9 | 437.92 | - |
| **PASEF** | 12 | 0.884 | 0.982 | 581.51 | 589.31 | - |
| **PASEF** | 12 | 0.982 | 1.17 | 757.88 | 769.91 | - |
| **PASEF** | 13 | 0.6 | 0.895 | 446.74 | 456.55 | - |
| **PASEF** | 13 | 0.895 | 0.99 | 596.31 | 604.3 | - |
| **PASEF** | 13 | 0.99 | 1.21 | 782.14 | 795.37 | - |
| **PASEF** | 14 | 0.6 | 0.903 | 464.4 | 473.26 | - |
| **PASEF** | 14 | 0.903 | 1 | 611.57 | 619.83 | - |
| **PASEF** | 14 | 1 | 1.25 | 808.15 | 821.93 | - |
| **PASEF** | 15 | 0.6 | 0.908 | 480.43 | 488.6 | - |
| **PASEF** | 15 | 0.908 | 1.015 | 627.34 | 635.85 | - |
| **PASEF** | 15 | 1.015 | 1.29 | 836.68 | 852.43 | - |
| **PASEF** | 16 | 0.6 | 0.914 | 495.43 | 503.27 | - |
| **PASEF** | 16 | 0.914 | 1.034 | 643.61 | 652.36 | - |
| **PASEF** | 16 | 1.034 | 1.33 | 869.21 | 886.99 | - |
| **PASEF** | 17 | 0.6 | 0.92 | 509.92 | 517.58 | - |
| **PASEF** | 17 | 0.92 | 1.054 | 660.35 | 669.33 | - |
| **PASEF** | 17 | 1.054 | 1.37 | 907.67 | 929.36 | - |
| **PASEF** | 18 | 0.6 | 0.934 | 524.18 | 531.79 | - |
| **PASEF** | 18 | 0.934 | 1.075 | 677.59 | 686.85 | - |
| **PASEF** | 18 | 1.075 | 1.41 | 955.76 | 983.17 | - |
| **PASEF** | 19 | 0.6 | 0.947 | 538.53 | 546.27 | - |
| **PASEF** | 19 | 0.947 | 1.095 | 695.62 | 705.39 | - |
| **PASEF** | 19 | 1.095 | 1.44 | 1021.11 | 1060.04 | - |
| **PASEF** | 20 | 0.6 | 0.96 | 552.79 | 560.3 | - |
| **PASEF** | 20 | 0.96 | 1.11 | 715.01 | 725.63 | - |
| **PASEF** | 20 | 1.11 | 1.45 | 1154.84 | 1250.64 | - |
